# PTH-Induced Behavioral and Metabolic Alterations in Mouse Models of Hyperparathyroidism

**DOI:** 10.1101/2025.10.07.681037

**Authors:** Lu Zhang, Yan Chen, Yuting Liu, Nian Liu, Yongliang Wu, Ze Du, Botai Li, Weikang Sun, Fan Yang

## Abstract

Parathyroid hormone (PTH) is a critical endocrine regulator of calcium homeostasis and bone remodeling and is widely employed in clinical settings through synthetic analogs for the treatment of osteoporosis. Although traditionally considered a peripheral regulator, emerging evidence indicates that PTH also exerts effects through the central nervous system (CNS). This study investigated the neuropsychiatric impact of elevated PTH and explored potential CNS involvement using multiple murine models of hyperparathyroidism (HPT). Distinct behavioral phenotypes were observed across models, indicating that psychiatric symptoms vary depending on disease etiology and progression. The effects of hPTH(1-34), a clinically approved PTH analog, were further assessed in male and female mice. Under pharmacological concentrations, hPTH(1-34) enhanced locomotor activity in males but induced mild anxiety-like behavior in females. These behavioral changes in females were independent of the estrus cycle and were amplified by ovariectomy. Metabolic analysis indicates PTH affects the basic metabolism by inhibiting the respiratory exchange ratio, promotes the energy expenditure and locomotion without affecting the food consumption in a 48hr range. To further investigate the molecular effect of PTH in the brain, a PTH1R-Cre mouse line was generated to map PTH receptor-1 (PTH1R) expression. Widespread expression of PTH1R was detected across the brain, including in both neuronal and non-neuronal cell populations. These findings suggest that PTH may influence behavior through interactions with PTH1R-expressing cells in brain vasculature and circumventricular regions. However, further studies are warranted to define the specific brain nuclei and cell types involved in PTH-driven modulation of neurobehavioral function.

## Introduction

Parathyroid hormone (PTH) plays a central role in maintaining calcium and phosphate balance through coordinated actions on bone, kidney, and intestine. In response to hypocalcemia, PTH is released and binds to the PTH receptor 1 (PTH1R), triggering calcium mobilization from bone, calcium reabsorption in the kidneys, and calcium absorption in the intestine. While the classical targets of PTH are well characterized, its actions in non-canonical tissues, particularly the central nervous system (CNS), remain poorly understood. Clinical observations have revealed that individuals with primary hyperparathyroidism (pHPT) frequently exhibit neuropsychiatric symptoms, including depression, anxiety, sleep disturbance, and cognitive impairment [1–4]. These symptoms often improve rapidly following parathyroidectomy, implicating a causal role for elevated PTH levels [5–11]. Similar psychiatric manifestations have also been reported in secondary hyperparathyroidism (sHPT), commonly associated with phosphate retention in chronic kidney disease, and may likewise be ameliorated through surgical intervention [12–17]. Collectively, these observations suggest that excess circulating PTH may contribute directly to neuropsychiatric dysfunction in both pHPT and sHPT patients, potentially through mechanisms involving CNS pathways that have yet to be fully delineated.

Early autoradiographic studies involving intracerebroventricular injection or direct incubation of radiolabeled PTH revealed specific binding within defined brain regions of rats and rabbits [18, 19], suggesting the presence of central PTH-responsive sites. Subsequent immunohistofluorescence studies confirmed the expression of PTH1R in the superior cervical ganglia of rats indicating PTH also targets the nervous system [20]. First identified in 1995 [21], PTH receptor 2 (PTH2R) exhibits distinct expression patterns across multiple brain regions. Functional investigations have implicated PTH2R in both acute and tonic nociceptive signaling, including inflammatory pain [22, 23], with disruption of PTH receptor-mediated pathways associated with increased expression of anxiety- and fear-related behaviors [24, 25]. Although PTH is capable of activating PTH2R, this receptor displays markedly greater affinity for tuberoinfundibular peptide of 39 residues (TIP39), a neuropeptide predominantly produced within the brain. However, the degree to which circulating PTH engages PTH2R under pathological conditions such as pHPT and sHPT has yet to be determined.

The ability of PTH to access the CNS across the blood-brain barrier (BBB) remains a subject of debate. Clinical studies in pediatric populations have reported detectable levels of PTH in the cerebrospinal fluid (CSF) of children at concentrations approximately 62.6% lower than levels in serum [26]. In contrast, another study involving individuals with Alzheimer’s disease found CSF PTH levels to be below the detection threshold of 3 ng/L [27]. Several investigations have concluded that neither peripheral infusion nor endogenous elevation of circulating PTH leads to a measurable increase in CSF PTH concentrations (reviewed by [28]). Based on transcriptomic data from the Allen Brain Atlas, the *Pth* gene is expressed in the mouse brain, suggesting the possibility of endogenous PTH secretion within the CNS. In patients with pHPT, elevated CSF PTH levels are associated with increased serum PTH, potentially reflecting BBB permeability changes [29]. Our previous study suggest that circulating PTH acts on the CNS via circumventricular organs (CVO), which bypass the BBB and permit direct hormone sensing [30], and contributing the regulatory effects of PTH on the CNS. Therefore, further study of the general expression of PTH receptors and the BBB permeability of PTH is required.

To address the above questions, this study compared behavioral outcomes in mouse models of PTH dysregulation and assessed metabolic and behavioral responses to pharmacological PTH exposure across sexes and physiological states. Our results indicated that PTH directly influences anxiety-like behavior and locomotor activity in a gender-dependent manner. The basic metabolic level of animals of different genders is also affected by PTH exposure in a dose-dependent manner. For both behavior and metabolic changes, female animals show higher tolerance to the impact of PTH. Given the widespread expression of PTH1R in central nervous system, the behavior changes are likely mediated through the cooperation of neuron and non-neuronal brain cell populations in CVOs. Collectively our findings provide important implications for clinical dosing, expected therapeutic efficacy, and potential adverse effects of PTH cross different genders, thus help clarify the broader physiological effects of PTH and the neuropsychiatric symptoms observed in PTH-related disorders.

## Results

### pHPT and sHPT induce distinct behavioral phenotypes in mice

Male PTH-Cre mice (Jackson Laboratory strain <u>005989</u>) were used to evaluate behavioral effects of endogenous PTH dysregulation. A subset of these animals spontaneously developed pHPT, characterized by parathyroid chief cell proliferation and elevated serum PTH concentrations (Fig. 1A, B). This phenotype did not appear in all animals and was not strictly age-dependent (Fig. 1C). Therefore, serum PTH levels were measured, and animals were stratified into low-PTH (<300 pg/mL) and high PTH (>300 pg/mL) groups for behavioral testing (Fig 1B). In the open-field test (OFT), high-PTH mice showed a marked increase in central zone activity, including a 366% increase in time spent in the central zone and a 282% increase in center entries, along with a 163% increase in total distance traveled (Fig 1D). In the elevated plus maze (EPM), no significant differences were observed between groups in open arm time or entries (Fig 1E). However, within the subgroup of animals with PTH levels exceeding 500 pg/mL, open arm time was positively correlated with serum PTH concentration (Fig 1E). In the tail suspension test (TST), no significant differences were detected in immobility time between the low-PTH and high-PTH groups (Fig 1F). From our studies, we find that spontaneous pHPT in male PTH-Cre mice is associated with increased exploratory behavior and locomotor activity, consistent with a hyperactivity phenotype. Notably, this contrasts with the typical age-related decline in activity often observed in male mice. However, the behavioral profile does not align with classical models of anxiety or depressive-like behavior, suggesting a distinct neurobehavioral signature linked to chronic PTH elevation.

**Figure 1.**
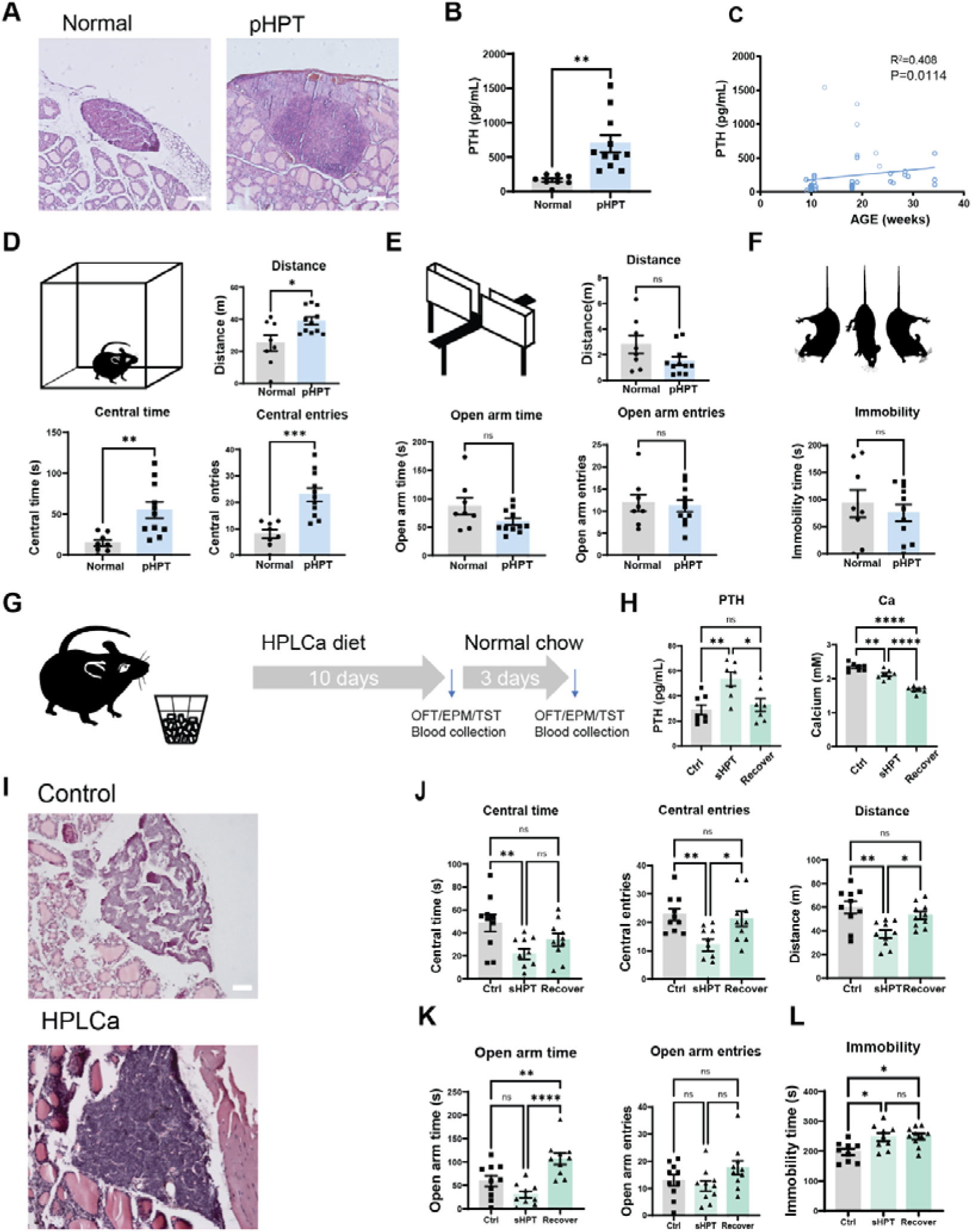
Behavioral alterations in mouse models of primary (pHPT) and secondary hyperparathyroidism (sHPT) (**A**–**E**) Behavioral and biochemical phenotypes in pHPT. **A**, Representative hematoxylin and eosin (H&E)-stained parathyroid glands from control and pHPT mice. Scale bar: 50 μm. **B**, Serum PTH concentrations in control and pHPT groups. N=8-11 **C**, Correlation between serum PTH levels and age across all groups. N=48 **D**, Total distance traveled (*upper*), time spent in the central zone (*lower left*), and number of central entries (*lower right*) in open-field test (OFT). N=8-11 **E**, Total distance traveled (*upper*), time spent in open arms (*lower left*), and number of open arm entries (*lower right*) in elevated plus maze (EPM). **F**, Immobility time in tail suspension test (TST). N=8-11 (**G**–**L**) Behavioral and biochemical phenotypes in diet-induced sHPT. N=8-11 **G**, Schematic overview of diet-induced sHPT model in mice. **H**, Serum PTH and total calcium levels in control, sHPT, and recovery groups. N=7-9 **I**, Representative H&E-stained sections of parathyroid glands from control and sHPT mice. Scale bar: 50 μm. **J**, Time spent in central zone (*left*), number of central entries (*middle*), and total distance traveled (*right*) in OFT. N=10 **K**, Time spent in open arms (*lower left*) and number of open arm entries (*right*) in EPM. N=10 **L**, Immobility time in TST. N=9-10 * P<0.05; ** P<0.01; *** P<0.005. All error bars and shaded areas show mean ± s.e.m.

To further investigate the mechanisms underlying psychiatric dysfunction in hyperparathyroidism, a dietary model of sHPT was established in male C57BL/6J mice. Eight-week-old mice were fed with a high-phosphate, low-calcium diet (HPLCa; 0.2% Ca, 1.2% P) for 10 days, followed by a 3-day recovery period on a standard diet containing 0.6% calcium and 0.6% phosphate (Fig 1G). Serum PTH concentrations increased by 86.2% after HPLCa feeding and returned to baseline following dietary normalization (Fig 1H, *left*). In contrast, serum calcium levels declined after HPLCa exposure and failed to recover after diet reversal (Fig 1H, *right*). Histological analysis revealed parathyroid chief cell proliferation in mice after 10 days of HPLCa treatment (Fig 1I). Behavioral assessment revealed a significant reduction in open-field activity during the sHPT phase, including a 55.3% decrease in time spent in the center, a 47.1% reduction in central entries, and a 37.3% decrease in total distance traveled. These deficits were reversed following dietary correction (Fig 1J). In the EPM test, sHPT animals exhibited a 48.6% decrease in open arm time, which also recovered with return to normal diet, while open arm entries remained unchanged throughout the experiment (Fig 1K). In the TST, immobility time—a proxy for depressive-like behavior— increased by 25.1% during sHPT and did not improve after dietary restoration (Fig 1L). These findings suggest that sHPT induces anxiety-like and depressive-like behaviors with partially reversible outcomes. This behavioral profile contrasts with the increased locomotor activity observed in pHPT models, reflecting distinct patterns associated with different forms of PTH dysregulation.

sHPT is often accompanied by systemic physiological alterations, including changes in serum calcium levels and renal dysfunction. To investigate whether kidney impairment could reproduce behavioral features observed in sHPT, a separate model of chronic kidney disease (CKD) associated sHPT was established using a high-phosphate, high-adenine diet (1.5% phosphate, 0.75% adenine) [31]. After 10 days of dietary exposure, mice exhibited reduced spontaneous activity in both the OFT and EPM (Fig S1). In the OFT, total distance travel declined by 49.9%, accompanied by a 67.3% reduction in central time and a 34.7% decrease in central entries. In the EPM, total distance was reduced by 22.5%. These findings indicate that both sHPT and the CKD associated sHPT models result in similar reductions in exploratory behavior, consistent with anxiety-like phenotypes. Given that sHPT frequently arises as a consequence of renal pathology in clinical settings, kidney dysfunction likely plays a primary role in driving the behavioral impairments observed in sHPT. In contrast, the increased locomotor activity seen in pHPT animals may reflect a direct effect of elevated circulating PTH.

### Peripheral administration of PTH induces sexually divergent behavioral responses

To examine the direct behavioral effect of circulating PTH, peripheral administration of recombinant human PTH(1–34)—the active fragment of the endogenous peptide—was evaluated in male and female C57BL/6J mice. Clinically, PTH receptor agonists such as teriparatide are widely used as anabolic treatments for osteoporosis in postmenopausal women and elderly men. In this study, hPTH(1–34) was administered at a clinically relevant concentration (0.33 μg/kg), and subsequent behavioral and metabolic outcomes were assessed.

At 30 min post-injection, serum PTH concentrations increased by 38.5% in males and 44.7% in females. However, no changes were detected in CSF in either sex (Fig 2A, B). Pharmacokinetic data for PTH(1-34) indicated rapid absorption and elimination, with peak serum concentration observed at 30 min following subcutaneous administration and undetectable levels by 3 h. Accordingly, behavioral testing was conducted during the 30–60-min window following peptide injection to capture the peak pharmacodynamic period.

**Figure 2.**
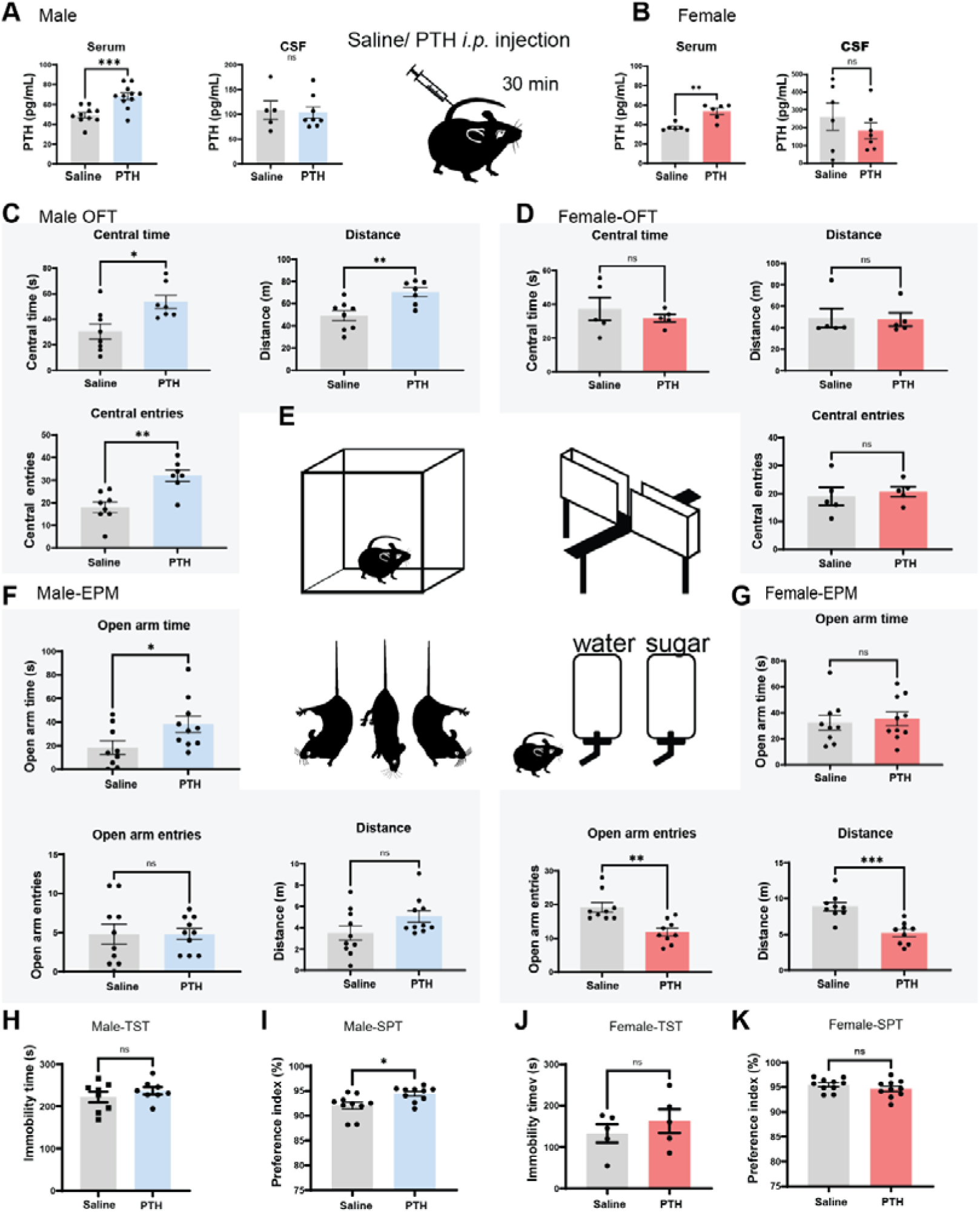
PTH administration alters behavior in male and female mice. (**A, B**) PTH concentrations in serum and cerebrospinal fluid (CSF) of male (A) and female (B) mice following saline or PTH administration. N=5-8. (**C**–**K**) Behavioral assessments of male and female mice after saline or PTH administration. N=5-10. (**C**, **D**) Time spent in central zone, number of central entries, and total distance traveled in male (C) and female (D) mice in OFT. (**E**) Schematic overview of behavioral assays, including the OFT, EPM, TST, and SPT. (**F**, **G**) Time spent in open arms and number of open arm entries in male (F) and female (G) mice in EPM. (**H**, **J**) Immobility time in male (H) and female (J) mice in TST. (**I**, **K**) Sucrose preference in male (I) and female (K) mice in SPT. * P<0.05; ** P<0.01; *** P<0.005. All error bars and shaded areas show mean ± s.e.m.

To evaluate the behavioral impact of circulating PTH, a series of assays reflecting anxiety-and depression-like phenotypes were conducted, including OFT, EPM, TST, and sucrose preference test (SPT). In male mice, administration of hPTH(1–34) significantly increased central zone time (76.1%), central entries (77.8%), and total distance traveled (43.0%) in the OFT compared to saline controls (Fig 2C). We also assessed the PTH impact on stress induced psychiatric disorder in male mice. It was found that PTH promotes the central traveling time, entrance and total traveling distance in the animals who went through chronic stress (Fig S2A). EPM testing also revealed a significant increase (107.4%) in time spent in the open arms following PTH injection (Fig 2F). However, this outcome differed from that of chronically stressed males, which showed a 43.1% reduction in open arm entries (Fig S2B). No significant changes were observed in TST immobility time in either unstressed or stressed males following PTH administration (Fig 2H, Fig S2C). Sucrose preference was modestly elevated (2.5%) in the PTH-treated non-stressed males (Fig 2I), with no effect observed in stressed counterparts (Fig S2D).

In contrast, behavioral responses to PTH were markedly different in female mice compared with male mice. In healthy adult females, PTH administration did not alter central time, central entries, or total distance traveled in the OFT (Fig 2D). However, in ovariectomized (OVX) females, PTH significantly reduced central time (44.0%) and central entries (36.4%) (Fig S2E). EPM analysis showed decreased open arm entries (38.2% in intact females and 48.4% in OVX females) and total distance traveled (41.3% and 39.3%, respectively) following PTH exposure (Fig 2G, Fig S2F). No significant differences were detected in TST immobility time or SPT outcomes in either intact or OVX females (Fig 2J, K; Fig S2G, H). To control for potential influence of the estrous cycle in females, behavioral assays were conducted across different estrous stages. No significant changes were observed in OFT, EPM, TST, or SPT parameters as a function of estrous status (Fig S2I–L), consistent with prior studies [32–34]. From this study, we could find that female animals are more resistant to the PTH exposure than male animals. The ablation of ovarian hormones has made the animals more vulnerable to the exogenous hormonal changes.

Aside from the psychiatric disorder associated behavior tests, we would further like to study the impact of PTH on the animals’ spontaneous behavior. This can be observed through a high-resolution 3D free-moving system. In this system, the animal body will be abstracted into a frame with 16 points. The movement of the frame will be categorized into 40 categories without supervision and then annotated and clustered manually[35]. In this study, we collected the 3D free-moving behavior of the animals with exposure to PTH or vehicles for 10 mins and analyzed the spontaneous behavior changes. The spontaneous behaviors were categorized into exploration, locomotion, and static states, encompassing subtypes such as up stretching, rearing, and sniffing (Fig 3A,B, Fig S3A). In male mice, PTH significantly reduced the proportion of time spent grooming, from 12.4% ± 6.5% to 6.9% ± 3.9% (Fig 3D). In females, only rearing behavior was modestly reduced, from 12.4% ± 2.8% to 9.5% ± 1.8% (Fig 3F). What is notable is that the ratio of spontaneous varies between male and female animals are also different: the grooming time of the male animals is significantly higher than female animals in both saline and PTH exposure group (Fig S3C). This difference is also observed in previous empathic behavior studies [36]. In our study, we found that male animals spontaneously have more grooming behavior than female animals. Kinetic analysis revealed a trend toward increased frame distance in males, although this did not reach statistical significance (Fig 3H). No such trend was observed in females (Fig 3J). Overall behavioral state distributions, including exploration and locomotion, were largely unaffected by PTH in both sexes (Fig S3B, C).

**Figure 3.**
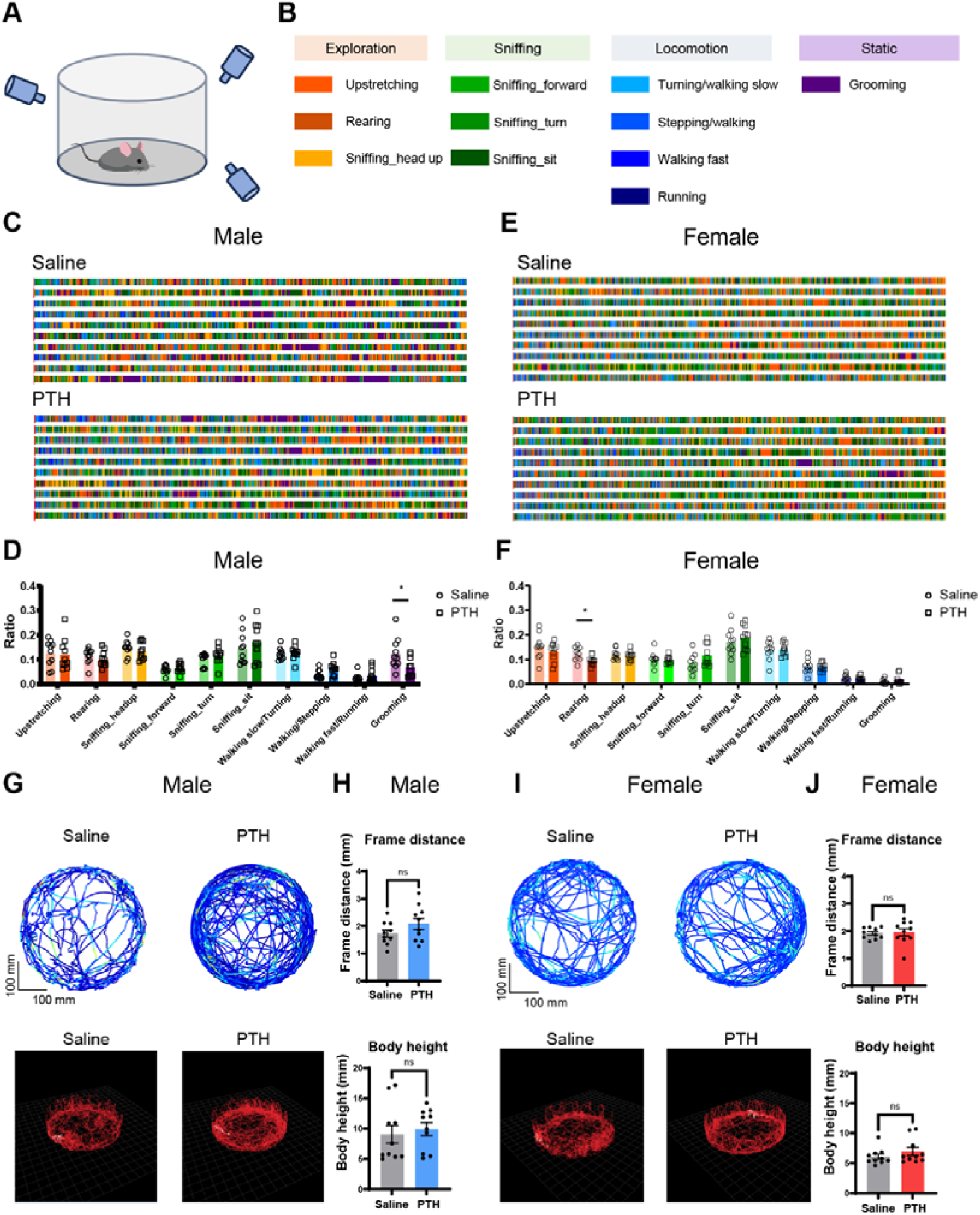
PTH-induced spontaneous behavioral changes captured under 3D supervision. **A**, Overview of 3D behavioral tracking system for monitoring spontaneous movement. **B**, Behavioral categories and annotation derived from unsupervised clustering. **C**, Ethograms of spontaneous behavior in male mice following saline or PTH administration. **D**, Statistic comparison of movement fractions of male mice under different annotated categories. **E**, Annotated ethogram of female mice under saline/PTH administration. **F**, Statistic comparison of movement fractions of female mice under different annotated categories. **G**–**J**, Movement dynamics of the 3D behavior of animals. **G**, Demonstration of animal behavior track in 2D (*upper*) and 3D (*lower*) of male animals after saline/PTH administration. **H**, Total frame distance of the male animals after saline/PTH injection (*upper*), body height of the male animals after saline/PTH injection (*lower*). **I**, Demonstration of animal behavior track in 2D (*upper*) and 3D (*lower*) of female animals after saline/PTH administration. **J**, Total frame distance of the female animals after saline/PTH injection (*upper*), body height of the male animals after saline/PTH injection (*lower*). N=10. * P<0.05. All error bars and shaded areas show mean ± s.e.m.

### PTH-induced metabolic changes in male and female mice

To determine whether the locomotor changes observed in the OFT and EPM tests following PTH administration were driven by systemic metabolic shifts or brain-induced behavioral change, metabolic profiling was conducted in male and female C57BL/6J mice using a metabolic cage system (Columbus, CLAMS-10M). Adult mice were habituated in metabolic cages for 48 h (Pretest phase), after which the animals received intraperitoneal (*i.p.*) injections of either hPTH(1–34) (0.33 mg/kg) or saline at the onset of Phase1, with continuous recording for 48 h. In Phase 2, the same mice received a higher dose of hPTH(1–34) (33 mg/kg) or saline, followed by an additional 48 h of monitoring in the cage. Each animal remained untreated during the Pretest phase and received only a single injection at the beginning of each subsequent phase. Several parameters indicated ongoing adaptation to the cage environment. For example, both respiratory exchange ratio (RER) and cumulative feed intake increased progressively across all phases, even in the saline-injected controls (Fig 4C– F), while Y-axis ambulatory values declined gradually over time (Fig 4G, H), suggesting habituation-related reductions. Therefore, comparisons focused on treatment differences between the saline- and PTH-treated groups, rather than changes across phases within one animal.

**Figure 4.**
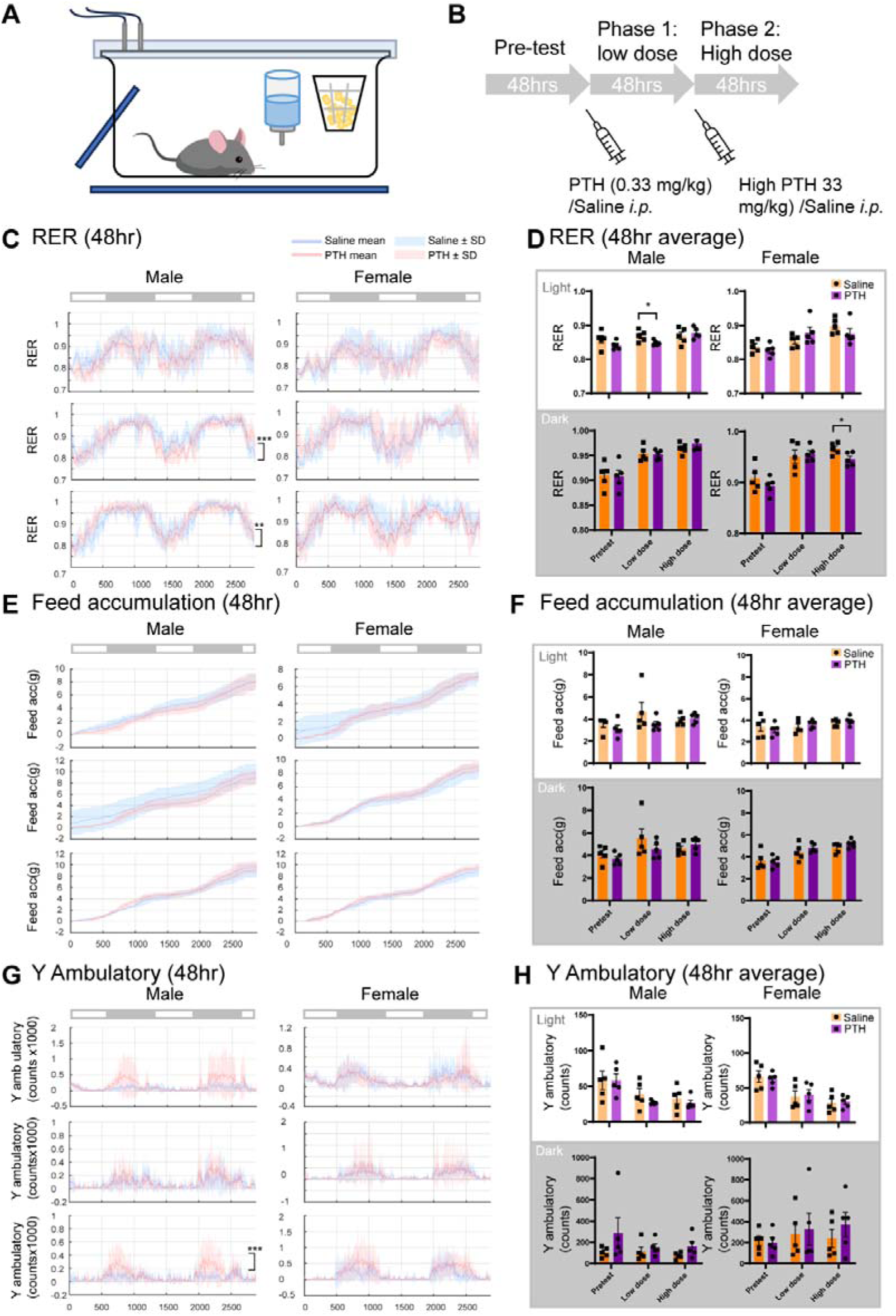
PTH-induced metabolic changes in mice. **A**, **B**, Schematic of metabolic phenotyping procedure in the Oxymax/CLAMS system. **C**, Respiration exchange ratio (RER) of male and female mice across experimental phases. Data are presented as mean ± SD. **D**, Quantification of 48-h average RER in male and female mice across phases. Data are presented as mean ± SEM. **E**, Cumulative food intake of male and female mice across phases. **F**, Quantification of 48-h average food intake in male and female mice across phases. Data are presented as mean ± SEM. **G**, Y-axis ambulatory activity of male and female mice across phases. Data are presented as mean ± SD. **H**, Quantification of 48-h average Y-axis ambulatory activity in male and female mice across phases. Data are presented as mean ± SEM. N=5. * P<0.05,** P<0.01,*** P<0.005. P value in **A**, **E**, **G** represents interaction effects analyzed by two-way ANOVA; P value in **D**, **F**, **H** represents significance analyzed by unpaired student’s t-tests.

Two-way ANOVA revealed that RER in male mice was significantly affected by both low- and high-dose PTH treatment (Fig 4C), while heat production in female mice was altered at the high dose (Fig S4A). Each 48-h phase was separated into light and dark periods, and average values were analyzed within each illumination condition. In males, RER was significantly reduced in the low-dose PTH group compared to saline controls during the light phase (Fig 4D, *upper left*). In females, RER was not affected until the high dose was administered (Fig 4D, *lower right*). These reductions suggest increased reliance on lipid oxidation for energy, consistent with previous findings showing that PTH promotes white adipose tissue browning [37]. No significant differences in cumulative food or water intake were observed in either sex across the 48-h period (Fig 4E, F; Fig S4C, D). Locomotor activity, as measured by X/Y-axis ambulation, showed no consistent treatment effect (Fig 4G, H; Fig E, F). Although two-way ANOVA indicated a significant difference in Y-axis ambulation in high-dose PTH-treated males (Fig 4G), this was not supported by unpaired *t-* tests conducted within light and dark phases (Fig 4H). These results indicate that PTH administration does not induce sustained changes in spontaneous locomotion within the 48-h metabolic cage assessment window. The behavioral effects observed in the OFT and EPM are therefore unlikely to be secondary to metabolic alterations and more likely reflect transient central mechanisms active during the acute phase of PTH signaling.

### Differential expression of PTH receptors in male and female mice

To investigate whether PTH-induced behavioral changes are primarily mediated through the CNS, brain expression patterns of PTH receptors were examined. Two PTH receptors respond to circulating PTH in mice: PTH1R and PTH2R. The EC_50_ of PTH1R (10^−10^ M) falls within the physiological serum PTH range (65–150 pg/mL). In contrast, PTH2R exhibits an EC_50_ for PTH of 10^−8^ M, approximately 100-fold higher [38, 39]. In this study, the PTH concentration in CSF was 0.011 ± 0.003 nM for males and 0.019 ± 0.012 nM for females following peripheral recombinant PTH administration, which is well below the activation threshold for PTH2R. Thus, the behavioral effects observed are unlikely to involve PTH2R signaling. we could be able to neglect the effect of PTH2R in this process.

To further examine sexually dimorphic PTH1R expression patterns, we constructed a PTH1r-cre animal. PTH1R-cre mice were crossed with Ai14 reporter mice to generate PTH1R-tdTomato mice (Fig S6). Whole-brain imaging of thin tissue sections revealed widespread tdTomato fluorescence, particularly in non-neuronal structures across various brain regions (Fig 5A). This finding is also supported by the MERFISH dataset on Allen Brain Atlas (Fig S5E, G) [40]. Transcriptomic data from the 10× scRNA-seq whole-brain atlas from the Brain Knowledge Platform confirmed broad expression of *Pth1r* across diverse cell types, including GABAergic neurons, glutamatergic neurons, and non-neuronal cell types such as astrocytes, oligodendrocytes, vascular cells, and leukocytes (Fig 5B, *left*; Fig S5A) [40]. In contrast, *Pth2r* expression was predominantly restricted to neurons (Fig 5B, *right*, Fig S5C).

**Figure 5.**
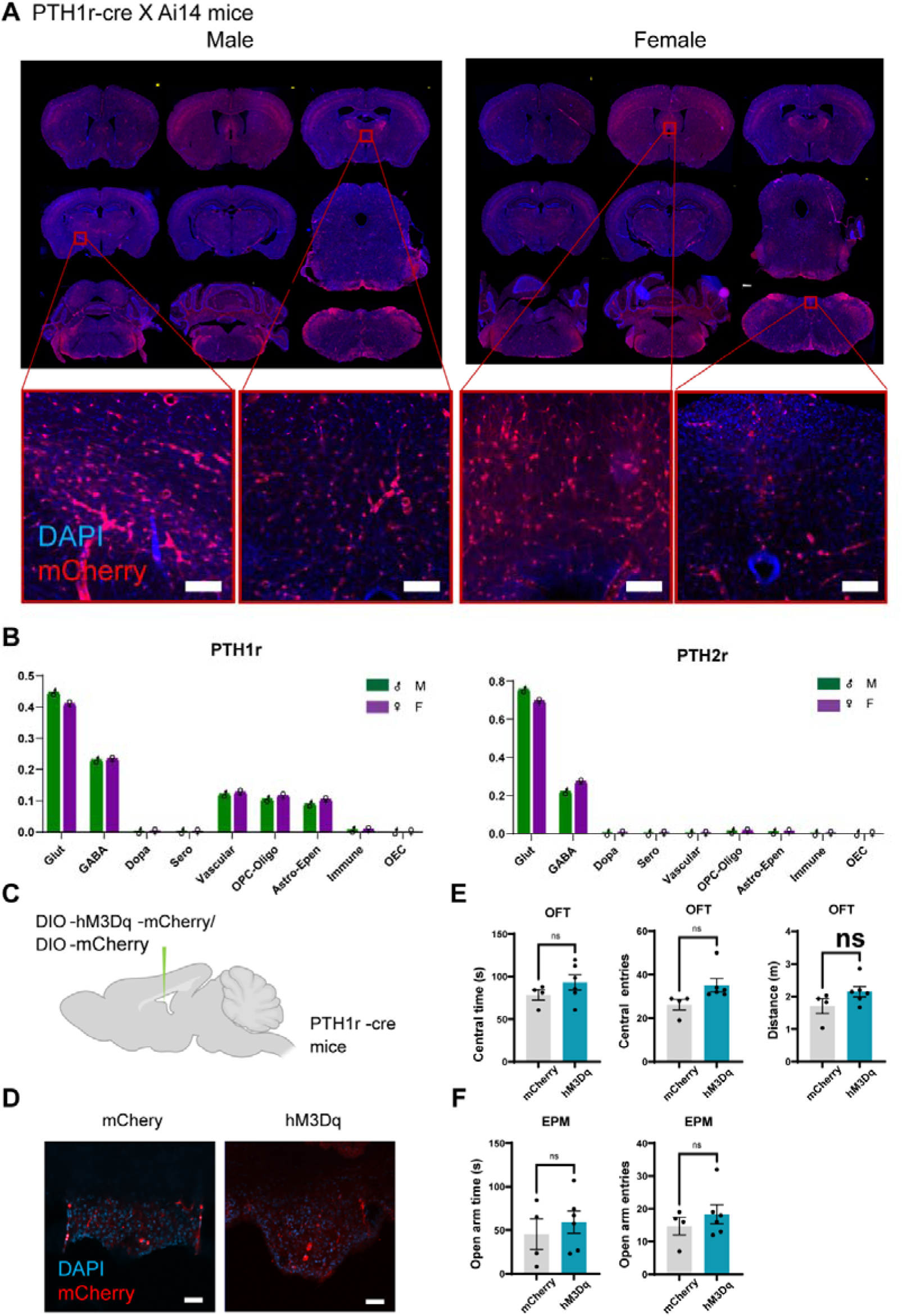
PTH1R expression and chemogenetic modulation in the brain. **A**, Representative whole-brain histological images showing PTH1R expression in male (*left*) and female (*right*) PTH1R-Cre × Ai14 mice. Scale bar: 100 μm. **B**, Cell-type distribution of PTH1R (*left*) and PTH2R (*right*) in male and female brains based on the from Allen Brain Atlas 10X scRNA-seq dataset. **C**, Schematic of chemogenetic manipulation targeting PTH1R-expressing cells in PTH1R-Cre mice. **D**, Representative image of AAV expression in the subfornical organ (SFO) of PTH1R-Cre mice. Scale bar: 50 μm. **E**, Quantification of time spent in central zone (*left*), number of central entries (*middle*), and total distance traveled (*right*) in the OFT under chemogenetic activation of SFO^PTH1R^ cells. N=4-6 **F**, Quantification of time spent in open arms (*left*) and number of open arm entries (*right*) in the EPM under chemogenetic regulation of SFO^PTH1R^ cells. Data are presented as mean ± SEM. N=4-6.

To further explore potential sex-based differences, we explored the proportions of *Pth1r*-expressing cell types were compared between male and female donors from the 10x scRNAseq whole brain dataset(Fig S5A,C). Although the proportion of the cells from female donors exhibited a slightly lower in glutamatergic neurons and a higher in non-neuronal cells than males, the differences were not statistically significant (Fig S5B). Spatially resolved analysis revealed that *Pth1r*-expressing glutamatergic neurons were enriched in the prelimbic/infralimbic/orbital cortex (PL-ILA-ORB), thalamus, and midbrain. In contrast, non-neuronal *Pth1r*-expressing cells dominated in the hippocampus and subamygdalar (sAMY) region, where GABAergic neurons also outnumbered glutamatergic neurons. In the hypothalamus, *Pth1r* expression was distributed relatively evenly across glutamatergic neurons, GABAergic neurons, and non-neuronal cells (Fig S5B). *Pth2r* expression patterns differed (Fig S5D), with glutamatergic neurons comprising over 80% of *Pth2r*-expressing cells in the PL-IL-ORB and thalamus. In the sAMY, hippocampus, and hypothalamus, GABAergic neurons were more prevalent among *Pth2r*-expressing cells. In the midbrain, glutamatergic and GABAergic neurons accounted for approximately 60% and 20% of *Pth2r* expressing cells, respectively. Overall, spatial heterogeneity in *Pth1r* and *Pth2r* expression outweighed sex-based differences in cell-type distribution.

According to both our findings in PTH1r-cre X Ai14 mice or the MERFISH data, PTH1r shows broadly expression on CVOs (Fig 5A, Fig S5F). To assess the functional contribution of PTH1R-expressing cells in CVOs, a chemogenetic approach was employed. A Cre-dependent activating viral vector was delivered to the subfornical organ (SFO) of PTH1R-Cre mice, which were then treated with the chemogenetic agonist. Activation of PTH1R-expressing cells in the SFO led to marginally increased center time, center entries, and total distance travel in the OFT (Fig 5E), as well as marginally increased open arm entries in the EPM (P=0.067, unpaired T-test, Fig.L5F, Supplementary table 1). These findings suggest that PTH1R-expressing cells in CVOs, particularly SFO, may contribute to the anxiolytic-like behavioral effects observed following peripheral PTH administration.

## Discussion

### Distinct behavioral changes in pHPT and sHPT

This study assessed anxiety- and depression-like behaviors in murine models of pHPT and sHPT. In the pHPT model, mice exhibited increased locomotor activity and central zone exploration in the OFT, despite being older than the controls. This pattern contrasts with the typical age-related decline in activity [41], suggesting that aging is unlikely to account for the behavioral changes observed in the high-dose PTH group. In contrast, sHPT mice displayed anxiety-like behavior during exposure to a HPLCa diet, which was reversed upon restoration of a normal chow diet. As serum calcium levels remained repressed following dietary normalization, recovery of exploratory behavior in the OFT and EPM was probably independent of calcium restoration. However, TST immobility time remained elevated during both the sHPT and recovery phases, suggesting a possible association with persistent calcium dysregulation. Additionally, mice subjected to a high-phosphate, high-adenine diet—a model of both sHPT and CKD—exhibited behavioral impairments similar to those observed in the HPLCa-induced sHPT model. These findings suggest that reduced central zone activity, diminished locomotion, and decreased open arm time are at least partially attributable to CKD-associated effects. Together, these results imply that the psychiatric manifestations observed in HPT patients may reflect the combined influence of elevated PTH, altered calcium homeostasis, and comorbid renal dysfunction.

### Male mice exhibit greater sensitivity to PTH-induced behavioral changes than females

This study demonstrated that peripheral administration of PTH in male mice elicited behavioral changes consistent with those observed in the pHPT model but opposite to the sHPT model in the OFT, supporting the hypothesis that anxiety-like behavior in sHPT are primarily driven by factors such as kidney dysfunction rather than PTH elevation alone. Notably, the behavioral effects of PTH differ between sexes and across test paradigms. In males, PTH exerted anxiolytic and modest antidepressant-like effects. To evaluate the consistency of these effects under chronic stress conditions, PTH was administered to stressed males. While anxiolytic behavior was retained in the OFT, anxiety-like behavior was paradoxically exacerbated in the EPM. Additionally, PTH had no significant effect on immobility time in the TST or sucrose preference in the SPT, indicating limited antidepressant efficacy at clinically relevant doses.

To examine the potential influence of gonadal hormones on PTH responsiveness in females, behavior tests were conducted 2 months post-ovariectomy. Following OVX, female mice exhibited increased susceptibility to PTH, displaying anxiety-like behavior in both the OFT and EPM. However, as in males, no significant changes were detected in TST or SPT outcomes. Overall, these findings indicate that male mice are more responsive to PTH-induced modulation of anxiety- and depression-like behaviors under normal conditions, while PTH sensitivity in females emerges following loss of ovarian hormones. This finding is also compliance to the clinical observation that female patients of pHPT at the age of primary menopause reports significant higher anxiety or depression scores than male [42, 43].

### PTH-induced metabolic change

Metabolic profiling over a 48-h period revealed that PTH promoted a shift in energy substrate utilization from carbohydrates to lipids, consistent with previous reports of PTH-induced adipose tissue browning [37]. Unlike the increased locomotor activity observed in the OFT, no consistent enhancement in locomotion was detected in the metabolic cage. Although some individuals exhibited elevated activity during the dark phase following PTH administration, the variability across animals obscured statistical significance. During the light phase, locomotor activity declined steadily across all treatment phases. As PTH was administered during the light phase, its effects on spontaneous locomotion may have been attenuated by circadian influences.

### Whole-brain expression of PTH1R in the mouse brain

Following peripheral administration of PTH, both serum and CSF concentrations fell within the effective activation range for PTH1R but remained well below the threshold required for PTH2R engagement. Notably, CSF PTH concentrations exceeded those in serum, a pattern that contradicts clinical findings [26, 27]. As PTH levels were assessed using enzyme-linked immunosorbent assay (ELISA) based on PTH-specific antibodies, it is possible that cross-reactivity with the PTH-related protein (PTHrP)—which is abundantly expressed in the CNS [44]—contributed to this discrepancy. Unexpectedly, PTH1R expression was detected broadly across the brain and across multiple cell types. While these results do not confirm CNS penetration of peripheral PTH under physiological conditions, the widespread expression of PTH1R in vascular cells suggests that PTH may act indirectly on the brain via vasculature and CVOs. Supporting this model, chemogenetic stimulation of PTH1R-expressing neurons in the SFO altered anxiety-like behaviors. However, the observed sex-dependent behavioral effects likely involve additional mechanisms, including receptor signaling dynamics, brain circuit modulation, and hormonal influences, which warrant further investigation.

In conclusion, this study systematically characterized anxiety- and depression-like behaviors in mouse models of pHPT and sHPT. The divergent behavioral phenotypes observed across models may help explain the psychiatric heterogeneity seen in HPT patients with differing etiologies and pathological stages. Furthermore, peripheral PTH administration elicited distinct behavioral responses in male and female mice, a difference that persisted after ovariectomy. The underlying mechanism driving this sexual divergence remains to be elucidated.

## Methods

### Animals

C57BL/6J mice (Beijing Vital River Laboratory Animal Technology Co., China), Ai14 reporter mice (Jackson Laboratory, 007908), and PTH1R-cre mice were maintained at the Shenzhen Institute of Advanced Technology, China. The PTH1R-Cre mice were generated on a C57BL/6J background by GemPharmatech Co., Ltd. via CRISPR-Cas9-mediated insertion of Cre recombinase downstream of the PTH1R promoter. All animals were housed under a specific pathogen-free (SPF) barrier facility in accordance with GB14925-2010 Laboratory Animals Requirement of Housing and Facilities. Mice were maintained in a temperature-controlled room (20–26 °C) on a 12-h light/dark cycle (07:00 lights on, 19:00 lights off), with free access to standard chow and water. All studies and experimental procedures were approved by the Institutional Animal Care and Use Committee (IACUC) of the Shenzhen Institute of Advanced Technology, Chinese Academy of Sciences.

### Secondary hyperparathyroidism (sHPT) models

sHPT was established in male C57BL/6J mice as described previously [45]. Following baseline behavioral testing under normal chow conditions, mice were fed a HPLCa diet (1.2% P, 0.2% Ca) for 10 days. Behavioral assays and blood collection were performed at the end of this period, after which mice were returned to normal chow for 3 days and reassessed. Parathyroid glands were collected postmortem and processed for hematoxylin and eosin (H&E) staining.

A chronic kidney disease dependent sHPT model was generated based on previously established protocols [46], with some modifications. Male C57BL/6J mice were randomly assigned to either a control group receiving normal chow or a treatment group (HP + Ad) fed a diet containing 1.5% P + 0.75% adenine (Beijing Keao Xieli Feed Co. Ltd., China).

### Viral injection and chemogenetic activation

Adult mice (6–8 weeks old) were anaesthetized under isoflurane anesthesia (3–4% induction, 1–2% maintenance in OL) and placed in a stereotaxic frame (RWD Life Science, Shenzhen, China). AAV9-DIO-mCherry or AAV9-DIO-hM3Dq-mCherry (1.5–2.5 E+12 PFU/mL) was loaded into a 5 μL-Hamilton syringe fitted with a 33-gauge needle. A total of 150 nL was injected at a rate of 100 nL/min into the subfornical organ (SFO coordinates: −0.58 mm AP, 0Lmm ML, −2.5 mm DV) based on the stereotaxic atlas. After surgery, mice were allowed at least 1 week to recover before behavioral experiments. For chemogenetic activation, clozapine-N-oxide (CNO; 1 mg/kg, *i.p.*; MCE, 34233-69-7) was administered 3 weeks post-virus injection, 30 min prior to behavioral testing. All AAV9 viral constructs were prepared in-house.

### Chronic stress model

Male C57BL/6J animals were restrain in pre-punched 50 mL centrifuge tubes, the restrain process lasts 2 hr daily for 10 days.

### Ovariectomy surgery

Bilateral ovariectomy (OVX) was performed on female C57BL/6J mice at 8 weeks under isoflurane anesthesia (3–4% induction, 1–2% maintenance in OL). After skin preparation and aseptic draping, small dorsolateral flank incisions (∼5 mm) were made to expose the peritoneal cavity. The ovarian fat pad was gently exteriorized, the oviduct and vascular pedicle were ligated with absorbable suture, and the ovary excised. The contralateral ovary was removed using the same procedure. Muscle and skin were closed with non-absorbable suture and tissue glue (Vetbond, 3M), respectively. After surgery, mice were allowed 8 week to recover before behavioral experiments.

### Histological analysis of brain and parathyroid gland (PTG)

Mice were deeply anaesthetized with sodium pentobarbital (100 mg/kg, *i.p.*) and perfused transcardially with phosphate-buffered saline (PBS) followed by 4% paraformaldehyde (PFA). Brains and PTGs were collected and fixed in 4% PFA at 4 °C for 48 h. Brain tissues were dehydrated in 30% sucrose solution for 72 h, while PTG samples were dehydrated through graded ethanol (70%–100%), cleared in xylene, and embedded in paraffin. Coronal brain sections (35 μm) and PTG sections (7 μm) were cut using a Cryostat microtome (Leica, CM1950) at −20 °C and room temperature microtome (Leica, RM2235), respectively. Brain cryosections were incubated with 0.5 mg/mL 4,6-diamidino-2-phenylindole dihydrochloride (DAPI; Thermo Fisher, D1306) for 1 min and mounted using Fluoromount-G (Southern Biotech, 0100-01). PTG sections were stained with H&E and mounted using Eukitt® Quick-hardening medium (Sigma, 03989). Images were acquired using Zeiss microscope AxioImagerZ2 or Olympus VS200-BU.

#### Fluorescence in situ Hybridization (FISH)

For FISH, anesthetized mice were intracardially perfused with DEPC-treated 1× PBS followed by 4% paraformaldehyde (containing DEPC) from Boster Biological Technology Co., Ltd. The brains were post-fixed in the same fixative for 24 hours at 4°C, then cryoprotected in 30% sucrose (in DEPC-treated PBS) until saturated. The samples were embedded in Tissue-Tek O.C.T. Compound (Sakura) and coronally sectioned at a thickness of 35 μm using a cryostat (Leica CM1950). A custom-designed probe targeting Pth1r mRNA was synthesized by Spatialfish Biotech Co., Ltd. The FISH procedure was carried out according to the manufacturer’s protocol. In brief, tissue sections were fixed, dehydrated, and then hybridized with the Pth1r probe in a humidified chamber overnight at 42°C. Following stringent washes, signal amplification was achieved via a rolling circle amplification (RCA) system using Phi29 DNA polymerase. Finally, the amplified signals were visualized by hybridization with fluorescently labeled probes. Sections were counterstained with DAPI and mounted with an anti-fade mounting medium. Multichannel fluorescence imaging was performed on a Leica STELLARIS 5 SR confocal microscope, with sequential acquisition of the FITC (Pth1r probe), tdTomato, and DAPI signals.

### Vaginal cytology

Mice were briefly restrained, and vaginal lavage was performed using 20 μL of PBS delivered through a fire-polished pipette tip, flushed gently 2–3 times to collect cells. Immediately after collection, the lavage fluid was expelled onto a microscope slide, air-dried, fixed, and subjected to H&E staining. Estrus cycle stages were determined based on the relative abundance and morphology of epithelial and leukocyte cell types. Images were acquired using Zeiss microscope AxioImagerZ2.

### Biochemical assessment

Serum PTH concentrations were assessed using a mouse PTH ELISA kit (LSBio, LS-5549). Total serum calcium levels were quantified using a colorimetric assay (Sigma, MAK022). All absorbance readings were acquired using a Nano Quant plate reader (Tecan, Infinite 200Pro).

### Behavioral tests

#### Elevated plus maze (EPM)

To assess locomotor activity and anxiety-like phenotypes subsequent to chronic disease or pharmacological interventions, mice were subjected to a single 5-min EPM trial, 30 min after *i.p.* administration of saline/PTH (0.33 μg/kg). conducted under treatment-blinded conditions.

Experimental mice were individually positioned on the central platform facing an open arm, with behavior recorded continuously via an overhead camera. Video recordings were automatically analyzed using ANY-maze™ software (Stoelting Co., Wood Dale, USA) to quantify time spent in open versus closed arms, total ambulatory distance traveled as a locomotor index, and frequency of open arm entries (defined by complete entry of all four paws). Anxiety-like behavior was operationally defined by a reduced ratio of time spent exploring open arms relative to total exploration time across all arms.

#### Open-field test (OFT)

The OFT was used to evaluate general locomotion and anxiety-like phenotypes over a 10-min period, 30 min after *i.p.* administration of saline/PTH (0.33 μg/kg), under experimenter-blinded conditions. Individual mice were positioned in the center of the arena, and behavior was recorded continuously throughout the session. ANY-maze™ (Stoelting Co., USA) was used to quantify time spent in the central and peripheral zones, total ambulatory distance traversed, and frequency of center entries (defined by complete entry of all four paws). Anxiety-like behavior was operationally defined by reduced proportional exploration of the central zone relative to total arena exploration time.

#### Tail suspension test (TST)

Behavioral despair was assessed using an automated TST system (BIO-TST, BioSeb, Chaville, France). Mice were suspended by the tail tip (adhesive tape applied <1 mm from the tip) from a hook positioned 20 cm above the testing platform. Each session lasted 6 min, during which immobility time was recorded as the primary outcome, modeling a condition analogous to human depression.

#### Sucrose preference test (SPT)

Anhedonia-like behavior was assessed using the SPT. Mice were individually housed and acclimated with two identical water bottles for 24 h, with food available *ad libitum*. On the second day, one bottle was replaced with a 1% sucrose solution. To eliminate potential spatial bias, bottle positions were switched on the third and fourth days. Fluid consumption was recorded over a 12-h testing period. The sucrose preference index (SPI) was calculated as: SPI (%) = [Volume of sucrose consumed (mL) / (Volume of sucrose consumed (mL) + Volume of water consumed (mL))] × 100%.

### Metabolism measurement

Metabolism assessment and analysis is performed by Columbus CLAMS-10M metabolism cage. To assess metabolic responses to PTH treatment—including oxygen consumption (VO_2_), carbon dioxide production (VCO_2_), food intake, water intake, respiratory exchange ratio (RER), energy expenditure, and locomotor activity—mice underwent comprehensive metabolic phenotyping. Following a 48-h acclimation period in an Oxymax/CLAMS-HC integrated metabolic monitoring system (Columbus Instruments), continuous data acquisition was conducted over the subsequent 48 h.

Prior to monitoring, body weight and baseline food mass per cage were recorded. Mice were individually housed in sealed Plexiglas chambers (40 × 25 × 20 cm) maintained at 22°C with a constant airflow of 0.5 L/min. Food and water were provided *ad libitum*. VO_2_ and VCO_2_ were sampled at 10-min intervals. RER was calculated as VCO_2_/VO_2_ to infer substrate utilization (1.0 = exclusive carbohydrate oxidation; 0.7 = exclusive fat oxidation). Energy expenditure is demonstrated as Heat (kcal/kg/hr) which was derived using the equation: Heat = (3.815+1.232×RER) × VO_2_/ body weight.

#### High-resolution 3D free-moving system and data analysis

The recording and analysis is constant to the previous paper[35]. Mice were individually placed in a transparent circular open field (50 cm diameter) and allowed to move freely. Four Intel RealSense D435 cameras were positioned at orthogonal sides to synchronously record 10 min of behavior. Multi-view recordings were processed using BehaviorAtlas Analyser software, following standardized and previously validated protocols. Sixteen skeletal key points were extracted per mouse to generate a 3D body model. A total of 39 kinematic parameters were computed to enable precise comparison of movement dynamics across experimental conditions. Behavioral frequency and sequence patterns were analyzed through 2D skeletal trajectories obtained from each camera using Deeplab Cut-trained models. Movement intensity (MI), scaled from 0 to 1 in arbitrary units, was visualized as heatmaps overlaid on skeletal projections. Behavior decomposition and unsupervised clustering were performed on 3D skeleton time-series data to identify movement clusters. The behavioral repertoire was defined by locomotor velocity and non-locomotor (NM) activity—such as grooming—that involves limb or organ movement without trunk displacement. A dynamic time alignment kernel was applied to quantify similarity between NM segments. High-dimensional NM features were embedded into the 2D NM space using Uniform Manifold Approximation and Projection (UMAP), which was combined with velocity to construct a 3D behavioral feature space. Unsupervised clustering was applied to classify distinct behavioral types. Supervised classification and dimensionality reduction enabled visual comparison of behavioral patterns across groups. Behavioral movement profiles—comprising 11 movement fractions and 116 transition types—were embedded into a 2D manifold using t-distributed stochastic neighbor embedding (t-SNE). A Gaussian kernel function was applied to this space to define support vector machine (SVM) decision boundaries for group classification.

#### Qualification and Statistical analysis

Statistical analyses were performed using GraphPad Prism (version 8.0) and Matlab (2025). Data collection and analysis were performed by investigators blinded to the conditions of the experiments. Experiments were randomized whenever possible. Sample sizes were not predetermined using statistical methods. The statistical tests used in this study include paired Student’s t-test, unpaired Student’s t-test, linear regression, and two-way analysis of variance. I The ‘*n*’ used for these analyses represents the number of mice, brain slices, or cells. All statistical details, including the statistical tests used, the exact value of ‘*n*’, etc., can be found in the figure legends. Data are reported as mean ± SEM or as a range from Min to Max; n.s. represents *p* > 0.05; ^∗^*p* < 0.05, ^∗∗^*p* < 0.01, ^∗∗∗^*p* < 0.001. A difference was accepted as statistically significant when probability (P) values were less than 0.05.

## Acknowledgements

This project was partly supported by the National Key R&D Program of China (2023YFA1801200 and 2023YFA1801202 F.Y.); Medical research Fund of Shenzhen Medical Academy of Research and Translation (C2301004) ; Shenzhen Science and Technology Research Funding (JCYJ20220818101414032).

## Author contribution

FY provided experimental resources and supervised this study. LZ designed and conducted the major experiments and wrote the manuscript; YC performed the FISH experiments, behavior experiments, and assisted manuscript writing; YL performed experiments of PTH induced anxiety like behavior in females, captured histological images and wrote the manuscript; NL performed experiments on behavior in pHPT animals; YW performed OFT and EPM experiments of PTH induced anxiety like behavior in OVX female animals; ZD assisted metabolic cage experiments; BL performed tissue preparation experiments; WS assisted the 3D behavior experiments.

## Supplementary Figures

**Supplementary Figure 1.**
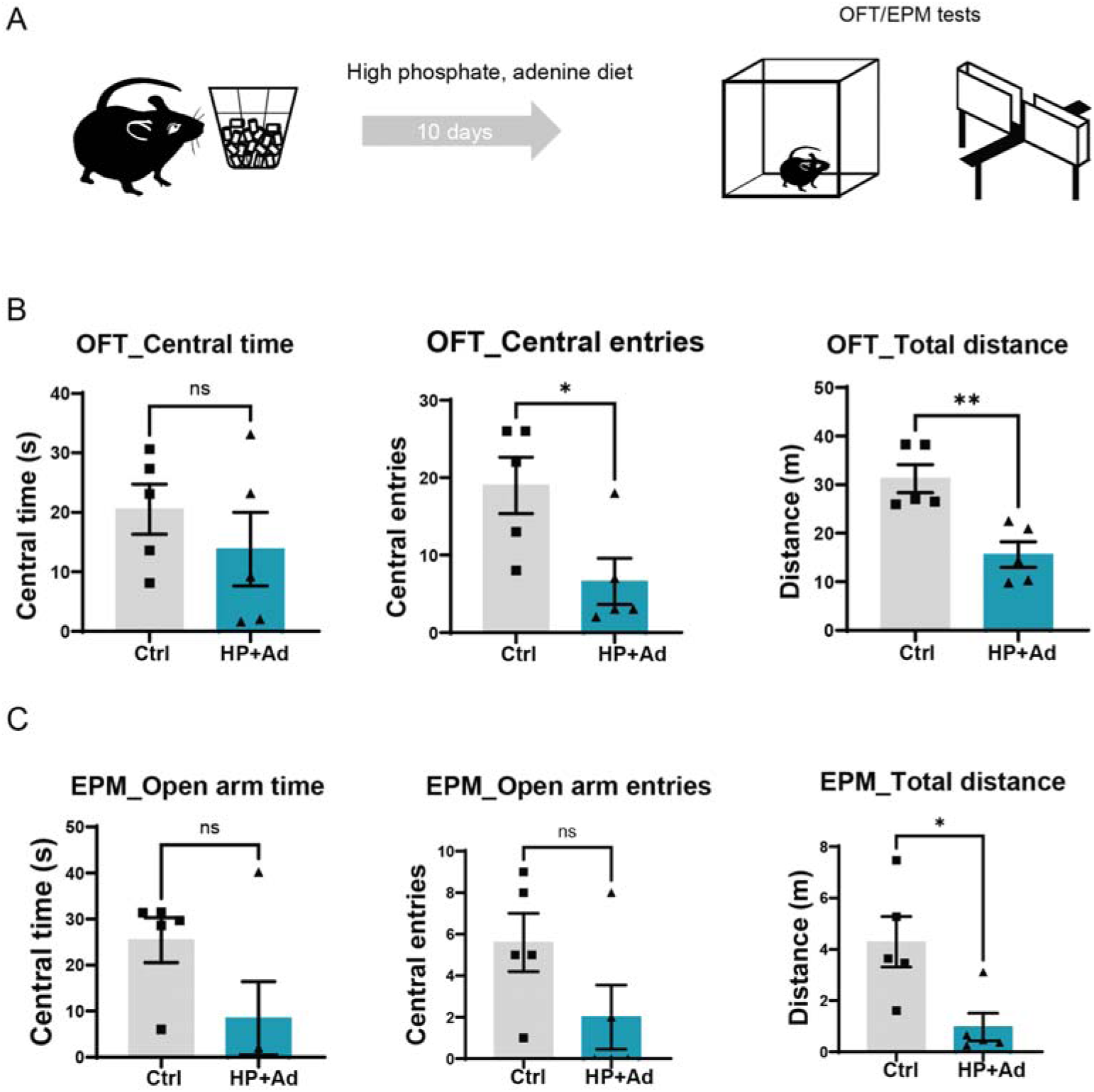
Behavioral alterations in mouse models of chronic kidney disease (CKD) (related to Figure 1) **A**, Schematic overview of diet-induced CKD model and behavioral assays. **B**, Time spent in central zone (*left*), number of central entries (*middle*), and total distance traveled (*right*) by CKD mice in the open-field test (OFT). N=5 **C**, Time spent in open arms (*left*), number of open arm entries (*middle*), and total distance traveled (*right*) by CKD mice in the elevated plus maze (EPM). N=5. * P<0.05; ** P<0.01; *** P<0.005. All error bars and shaded areas show mean ± s.e.m.

**Supplementary Figure 2.**
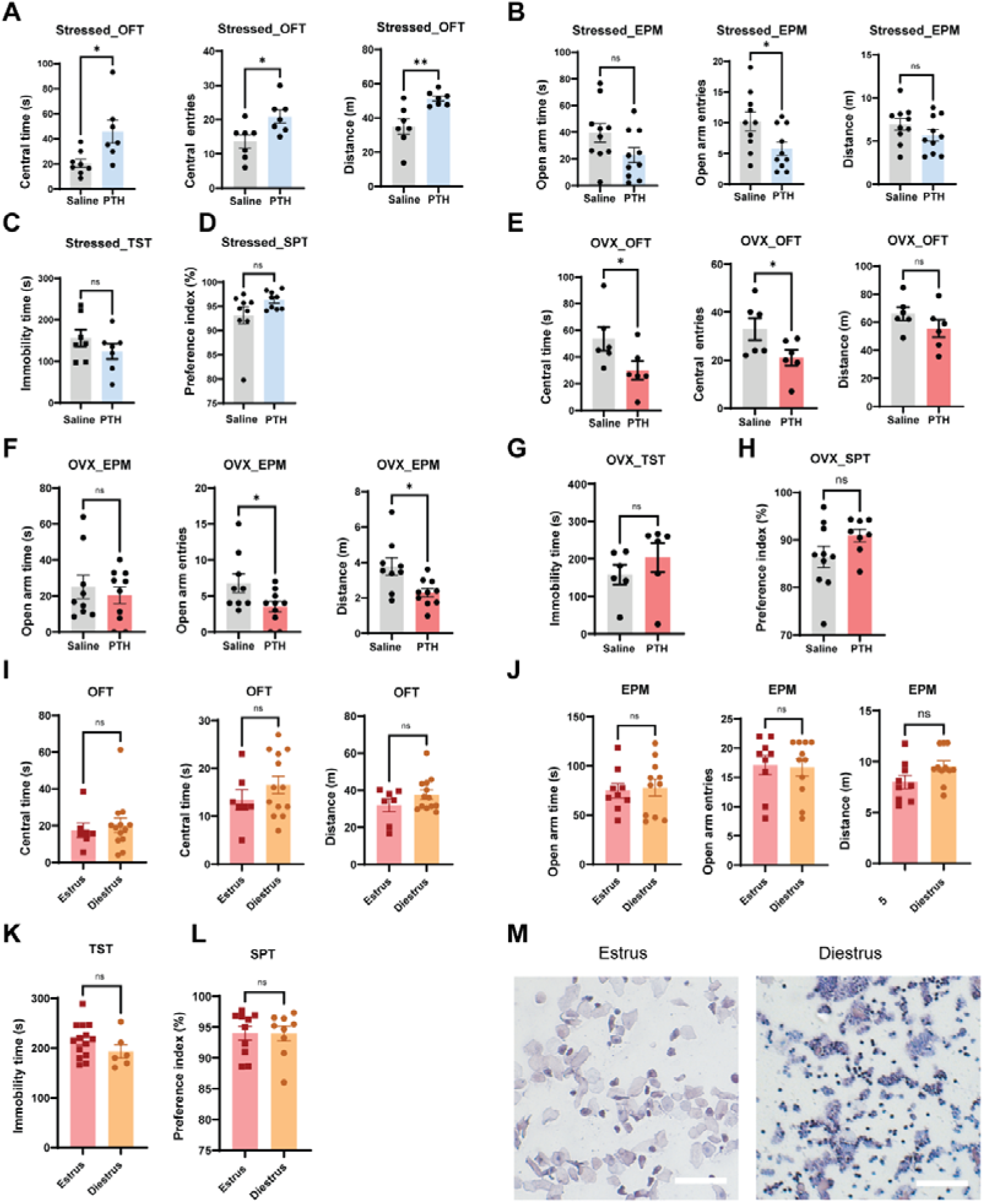
Behavioral effects of PTH in male and female mice (related to Figure 2) **A**, Time spent in central zone (*left*), number of central entries (*middle*), and total distance traveled (*left*) by chronically stressed male mice in OFT. N=7-8 **B**, Time spent in open arms (*left*), number of open arm entries (*middle*), and total distance traveled (*right*) by chronically stressed male mice in EPM. N=10 **C**, Immobility time of chronically stressed male mice in TST. N=7 **D**, Sucrose preference index of chronically stressed male mice in SPT. N=9 **E**, Time spent in central zone (*left*), number of central entries (*middle*), and total distance traveled (*left*) by ovariectomized female mice in OFT. N=6 **F**, Time spent in open arms (*left*), number of open arm entries (*middle*), and total distance traveled (*right*) by ovariectomized female mice in EPM. N=9-10 **G**, Immobility time of ovariectomized female mice in TST. N=6 **H**, Sucrose preference index of ovariectomized female mice in SPT. N=9-10 **I**, Time spent in central zone (*left*), number of central entries (*middle*), and total distance traveled (*left*) by female mice in different estrus phases in OFT. N=7-13 **J**, Time spent in open arms (*left*), number of open arm entries (*middle*), and total distance traveled (*right*) by female mice in different estrus phase in EPM. N=9-11 **K**, Immobility time of female mice in different estrus phase in TST. N=6-14 **L**, Sucrose preference index of female mice in different estrus phase in SPT. N=9-10 **M**, Representative vaginal cytology images from female mice in estrus and diestrus stages. Scale bar: 100 μm.

**Supplementary Figure 3.**
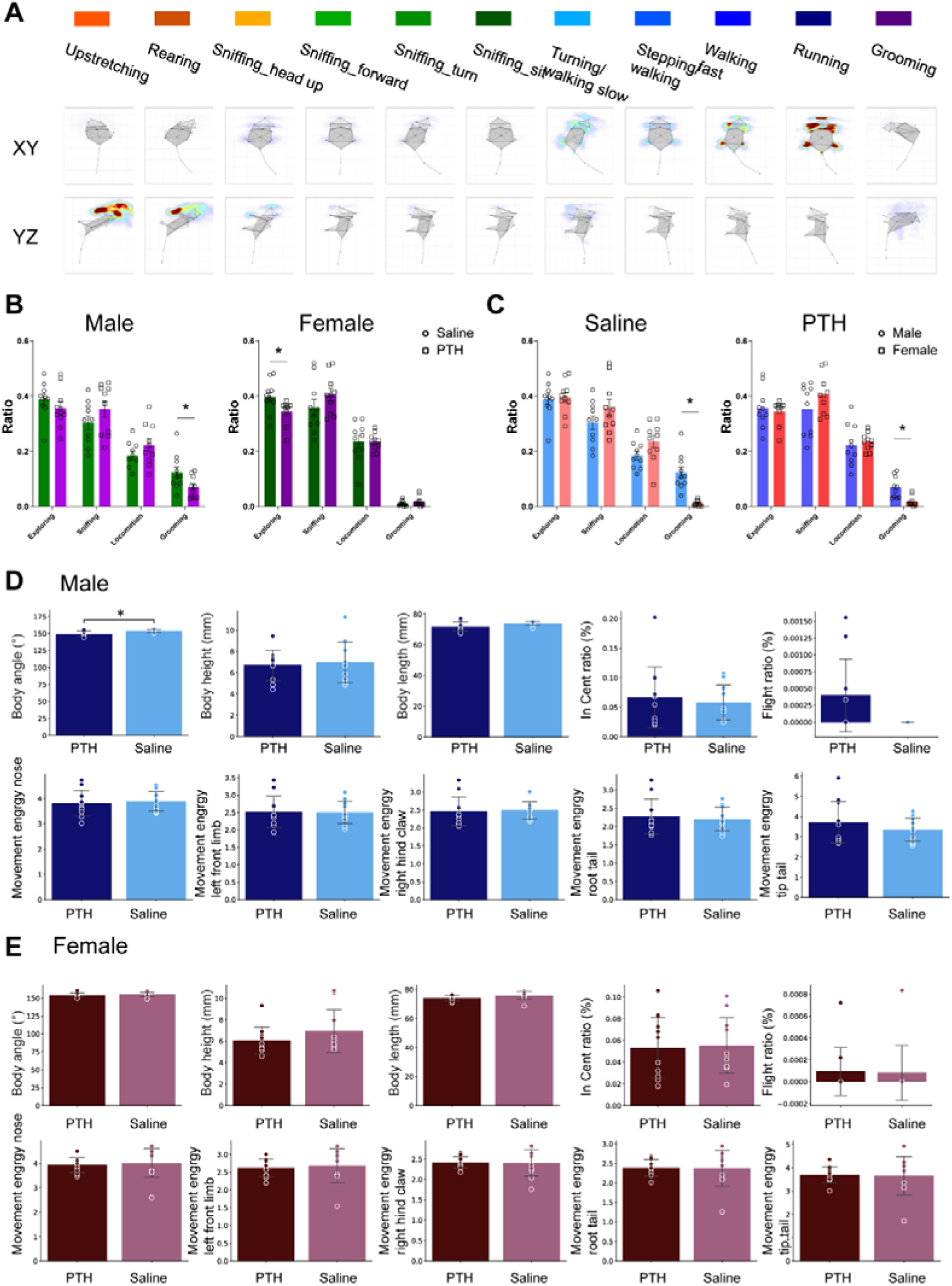
PTH-induced spontaneous behavioral changes captured under 3D supervision (related to Figure 3) **A**, Representative body posture and speed profiles for each behavioral category in ethogram. **B**, Proportional changes in clustered movement categories in male and female mice following PTH administration comparing between genders. **C**, Proportional changes in clustered movement categories in male and female mice following PTH administration comparing between treatments. (**D**, **E**) PTH-induced changes in dynamic movement parameters in male (**D**) and female (**E**) mice, including body angle, body height, body length, in-center ratio, flight ratio, movement energy of the nose, left forelimb, tail root, and tail tip. N=10. * P<0.05. All error bars and shaded areas show mean ± s.e.m.

**Supplementary Figure 4.**
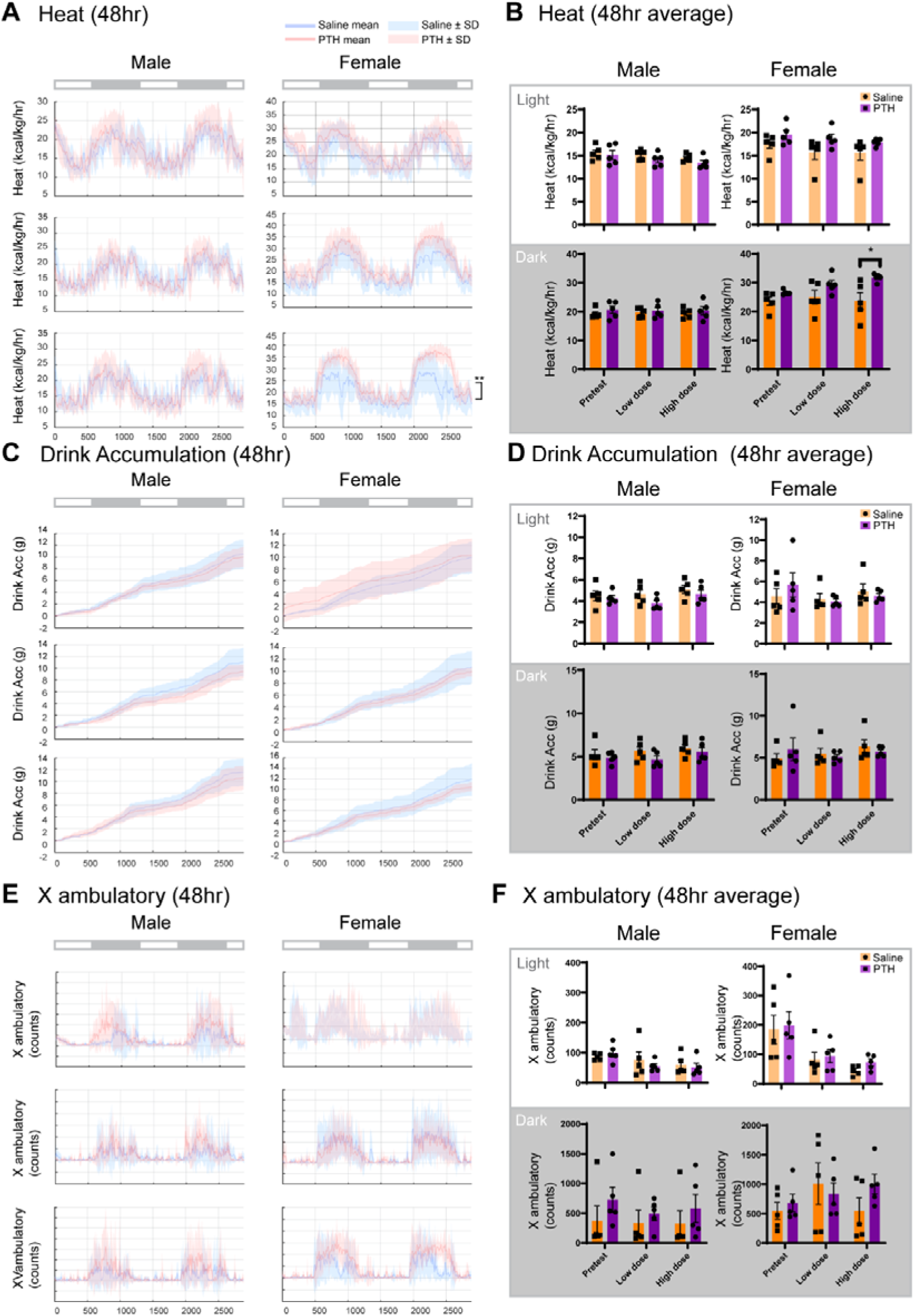
PTH-induced metabolic changes in mice (related to Figure 4) **A**, Heat production of male and female mice across experimental phases. Data are presented as mean ± SD. **B**, Quantification of 48-h average heat production in male and female mice across phases. Data are presented as mean ± SEM. **C**, Cumulative water intake in male and female mice across phases. Data are presented as mean ± SD. **D**, Quantification of 48-h average water intake in male and female mice across phases. Data are presented as mean ± SEM. **E**, X-axis ambulatory activity in male and female mice across phases. Data are presented as mean ± SD. **F**, Quantification of 48-h average X-axis ambulatory activity in male and female mice across phases. Data are presented as mean ± SEM. N=5. * P<0.05,** P<0.01,*** P<0.005. P value in **A**, **C**, **E** represents interaction effects analyzed by two-way ANOVA; P value in **B**, **D**, **F** represents significance analyzed by unpaired student’s t-tests.

**Supplementary Figure 5.**
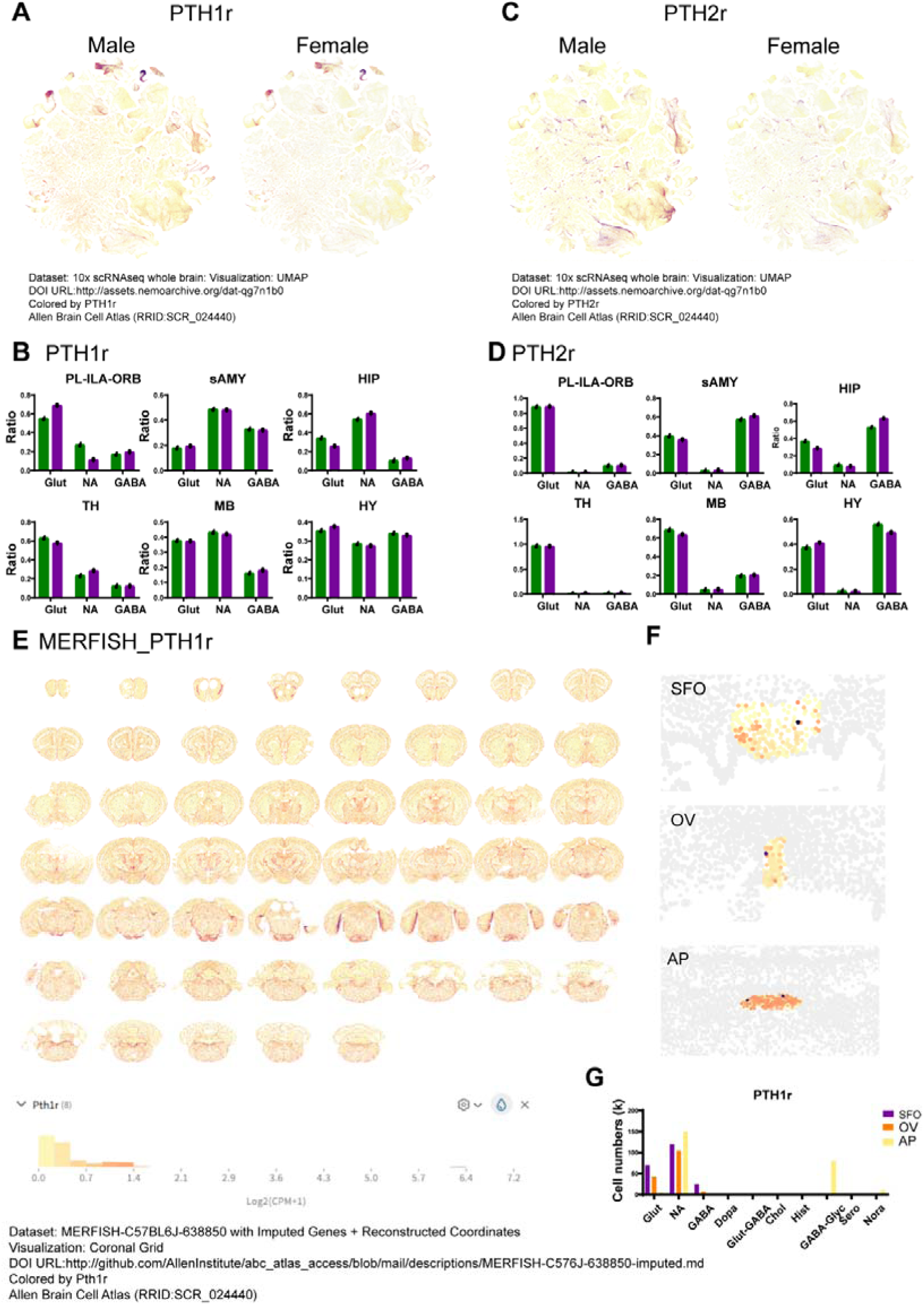
Expression of PTH1R and PTH2R in mouse brain from the Allen Brain Atlas (related to Figure 5) (**A–D**) Expression of *Pth1r* and *Pth2r* in the Allen Brain Atlas 10X scRNA-seq whole-brain dataset. **A**, UMAP visualization of *Pth1r* in male (*left*) and female (*right*) brains. **B**, Proportion of glutamatergic, GABAergic, and non-neuronal cells among *Pth1r*-expressing cells across spatial regions in male and female donors. **C**, UMAP visualization of *Pth2r* in male (*left*) and female (*right*) brains. **D**, Proportion of glutamatergic, GABAergic, and non-neuronal cells among *Pth2r*-expressing cells across spatial regions in male and female donors. (**E–H**) *Pth1r* expression in MERFISH-C57BL6J whole-brain dataset. **E**, Whole-brain distribution of *Pth1r*-expressing cells colored by expression intensity. **F**, Distribution of *Pth1r*-expressing cells colored by expression intensity in subfornical organ (SFO), organum vasculosum (OV), and area postrema (AP). **G**, Distribution of different neurotransmitters in *Pth1r*-expressing cells in the SFO, OV, and AP.

**Supplementary Figure 6.**
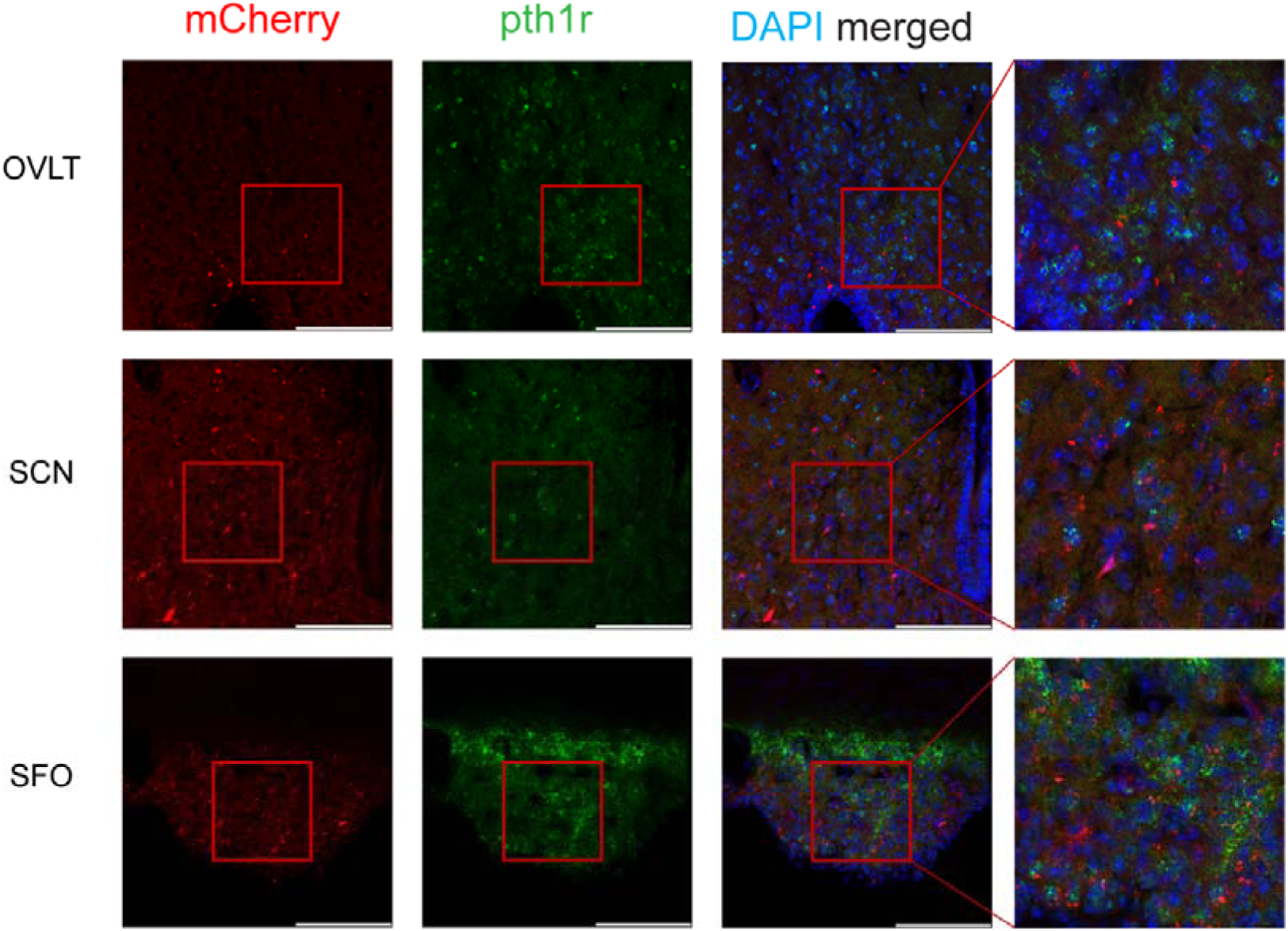
Colocalization of *in situ* mRNA hybridization staining of *pth1r* with tdTomato signals in PTH1R X Ai14 mice brain (related to Figure 5) *In situ* hybridization staining of the mRNA of PTH1R in SFO, OVLT and SCN in PTH1R X Ai14 mice. Scale bar, 100 μm.

**Supplementary Table 1.** Statistics and *P*-values related to Figures 1–5.

| Figure | Title | Compare groups | *P* value |  |
| --- | --- | --- | --- | --- |
| Figure 1B | PTH | Normal vs. pHPT | 0.0024 | Unpaired *t*-test |
| Figure 1C |  | Age vs. PTH | 0.0114 |  |
| Figure 1D | Distance | Normal vs. pHPT | 0.034 | Unpaired *t*-test |
|  | Central time | Normal vs. pHPT | 0.0036 | Unpaired *t*-test |
|  | Central entries | Normal vs. pHPT | 0.0004 | Unpaired *t*-test |
| Figure 1E | Distance | Normal vs. pHPT | 0.084 | Unpaired *t*-test |
|  | Open arm time | Normal vs. pHPT | 0.082 | Unpaired *t*-test |
|  | Open arm entries | Normal vs. pHPT | 0.76 | Unpaired *t*-test |
| Figure 1F | Immobility | Normal vs. pHPT | 0.56 | Unpaired *t*-test |
| Figure 1H | PTH | Normal vs. sHPT | 0.0023 | Unpaired *t*-test |
|  |  | sHPT vs. Recover | 0.017 | Unpaired *t*-test |
|  | Ca | Normal vs. sHPT | 0.0055 | Unpaired *t*-test |
|  |  | sHPT vs. Recover | <0.001 | Unpaired *t*-test |
| Figure 1J | Central time | Normal vs. sHPT | 0.0064 | Unpaired *t*-test |
|  |  | sHPT vs. Recover | 0.0097 | Unpaired *t*-test |
|  | Central entries | Normal vs. sHPT | 0.0017 | Unpaired *t*-test |
|  |  | sHPT vs. Recover | 0.014 | Unpaired *t*-test |
|  | Distance | Normal vs. sHPT | 0.009 | Unpaired *t*-test |
|  |  | sHPT vs. Recover | 0.032 | Unpaired *t*-test |
| Figure 1K | Open arm time | Normal vs. sHPT | 0.044 | Unpaired *t*-test |
|  |  | sHPT vs. Recover | <0.001 | Unpaired *t*-test |
| Figure 1K | Open arm entries | Normal vs. sHPT | 0.489 | Unpaired *t*-test |
|  |  | sHPT vs. Recover | 0.057 | Unpaired *t*-test |
| Figure 1L | Immobility | Normal vs. sHPT | 0.012 | Unpaired *t*-test |
|  |  | sHPT vs. Recover | 0.896 | Unpaired *t*-test |
| Figure 2A | Serum | Saline vs. PTH | 0.0004 | Unpaired *t*-test |
|  | CSF | Saline vs. PTH | 0.834 | Unpaired *t*-test |
| Figure 2B | Serum | Saline vs. PTH | 0.0011 | Unpaired *t*-test |
|  | CSF | Saline vs. PTH | 0.370 | Unpaired *t*-test |
| Figure 2C | Central time | Saline vs. PTH | 0.0130 | Unpaired *t*-test |
|  | Central entries | Saline vs. PTH | 0.0016 | Unpaired *t*-test |
|  | Distance | Saline vs. PTH | 0.0047 | Unpaired *t*-test |
| Figure 2D | Central time | Saline vs. PTH | 0.473 | Unpaired *t*-test |
|  | Central entries | Saline vs. PTH | 0.663 | Unpaired *t*-test |
|  | Distance | Saline vs. PTH | 0.905 | Unpaired *t*-test |
| Figure 2F | Open arm time | Saline vs. PTH | 0.042 | Unpaired *t*-test |
|  | Open arm entries | Saline vs. PTH | 1 | Unpaired *t*-test |
|  | Distance | Saline vs. PTH | 0.093 | Unpaired *t*-test |
| Figure 2G | Open arm time | Saline vs. PTH | 0.001 | Unpaired *t*-test |
|  | Open arm entries | Saline vs. PTH | 0.581 | Unpaired *t*-test |
|  | Distance | Saline vs. PTH | 0.0002 | Unpaired *t*-test |
| Figure 2H | TST | Saline vs. PTH | 0.347 | Unpaired *t*-test |
| Figure 2I | SPT | Saline vs. PTH | 0.0133 | Unpaired *t*-test |
| Figure 2J | TST | Saline vs. PTH | 0.439 | Unpaired *t*-test |
| Figure 2K | SPT | Saline vs. PTH | 0.250 | Unpaired *t*-test |
| Figure 3D | Up stretching | Saline vs. PTH | 0.838222757 | Unpaired *t*-test |
|  | Rearing | Saline vs. PTH | 0.38496167 | Unpaired *t*-test |
|  | Sniffing_headup | Saline vs. PTH | 0.355463237 | Unpaired *t*-test |
|  | Sniffing_forward | Saline vs. PTH | 0.442586184 | Unpaired *t*-test |
|  | Sniffing_turn | Saline vs. PTH | 0.229396804 | Unpaired *t*-test |
|  | Sniffing_sit | Saline vs. PTH | 0.42936853 | Unpaired *t*-test |
|  | Walking slow/Turning | Saline vs. PTH | 0.704345738 | Unpaired *t*-test |
|  | Walking/Stepping | Saline vs. PTH | 0.057539694 | Unpaired *t*-test |
|  | Walking fast/Running | Saline vs. PTH | 0.318017054 | Unpaired *t*-test |
|  | Grooming | Saline vs. PTH | 0.034874736 | Unpaired *t*-test |
| Figure 3F | Up stretching | Saline vs. PTH | 0.31864334 | Unpaired *t*-test |
|  | Rearing | Saline vs. PTH | 0.013120115 | Unpaired *t*-test |
|  | Sniffing_headup | Saline vs. PTH | 0.650209549 | Unpaired *t*-test |
|  | Sniffing_forward | Saline vs. PTH | 0.677613538 | Unpaired *t*-test |
|  | Sniffing_turn | Saline vs. PTH | 0.089112935 | Unpaired *t*-test |
|  | Sniffing_sit | Saline vs. PTH | 0.416410934 | Unpaired *t*-test |
|  | Walking slow/Turning | Saline vs. PTH | 0.844178217 | Unpaired *t*-test |
|  | Walking/Stepping | Saline vs. PTH | 0.872226836 | Unpaired *t*-test |
|  | Walking fast/Running | Saline vs. PTH | 0.687653934 | Unpaired *t*-test |
|  | Grooming | Saline vs. PTH | 0.284878136 | Unpaired *t*-test |
| Figure 3H | Frame distance | Saline vs. PTH | 0.151 | Unpaired *t*-test |
|  | Body height | Saline vs. PTH | 0.630 | Unpaired *t*-test |
| Figure 3J | Frame distance | Saline vs. PTH | 0.820 | Unpaired *t*-test |
|  | Body height | Saline vs. PTH | 0.284 | Unpaired *t*-test |
| Figure 4C | Male-Pretest | Saline vs. PTH | 0.357481683 | Two-way ANOVA, interaction |
|  | Male-Phase1 | Saline vs. PTH | 9.55718E-11 | Two-way ANOVA, interaction |
|  | Male-Phase2 | Saline vs. PTH | 0.008811874 | Two-way ANOVA, interaction |
|  | Female-Pretest | Saline vs. PTH | 1 | Two-way ANOVA, interaction |
|  | Female-Phase1 | Saline vs. PTH | 0.999916186 | Two-way ANOVA, interaction |
|  | Female-Phase2 | Saline vs. PTH | 0.985140757 | Two-way ANOVA, interaction |
| Figure 4D | Male-light-Pretest | Saline vs. PTH | 0.175541 | Unpaired *t*-test |
|  | Male-light-Low dose | Saline vs. PTH | 0.039302 | Unpaired *t*-test |
|  | Male-light-High dose | Saline vs. PTH | 0.55566 | Unpaired *t*-test |
|  | Female-light-Pretest | Saline vs. PTH | 0.175541 | Unpaired *t*-test |
|  | Female-light-Low dose | Saline vs. PTH | 0.039302 | Unpaired *t*-test |
|  | Female-light-High dose | Saline vs. PTH | 0.55566 | Unpaired *t*-test |
|  | Male-dark-Pretest | Saline vs. PTH | 0.908077 | Unpaired *t*-test |
|  | Male-dark-Low dose | Saline vs. PTH | 0.862571 | Unpaired *t*-test |
|  | Male-dark-High dose | Saline vs. PTH | 0.341816 | Unpaired *t*-test |
|  | Female-dark-Pretest | Saline vs. PTH | 0.313991 | Unpaired *t*-test |
|  | Female-dark-Low dose | Saline vs. PTH | 0.722794 | Unpaired *t*-test |
|  | Female-dark-High dose | Saline vs. PTH | 0.030336 | Unpaired *t*-test |
| Figure 4E | Male-Pretest | Saline vs. PTH | 1 | Two-way ANOVA, interaction |
|  | Male-Phase1 | Saline vs. PTH | 1 | Two-way ANOVA, interaction |
|  | Male-Phase2 | Saline vs. PTH | 1 | Two-way ANOVA, interaction |
|  | Female-Pretest | Saline vs. PTH | 1 | Two-way ANOVA, interaction |
|  | Female-Phase1 | Saline vs. PTH | 1 | Two-way ANOVA, interaction |
|  | Female-Phase2 | Saline vs. PTH | 0.759662435 | Two-way ANOVA, interaction |
| Figure 4F | Male-light-Pretest | Saline vs. PTH | 0.366361 | Unpaired *t*-test |
|  | Male-light-Low dose | Saline vs. PTH | 0.244213 | Unpaired *t*-test |
|  | Male-light-High dose | Saline vs. PTH | 0.567175 | Unpaired *t*-test |
|  | Female-light-Pretest | Saline vs. PTH | 0.59388 | Unpaired *t*-test |
|  | Female-light-Low dose | Saline vs. PTH | 0.461529 | Unpaired *t*-test |
|  | Female-light-High dose | Saline vs. PTH | 0.462337 | Unpaired *t*-test |
|  | Male-dark-Pretest | Saline vs. PTH | 0.317251 | Unpaired *t*-test |
|  | Male-dark-Low dose | Saline vs. PTH | 0.321352 | Unpaired *t*-test |
|  | Male-dark-High dose | Saline vs. PTH | 0.298944 | Unpaired *t*-test |
|  | Female-dark-Pretest | Saline vs. PTH | 0.838204 | Unpaired *t*-test |
|  | Female-dark-Low dose | Saline vs. PTH | 0.372698 | Unpaired *t*-test |
|  | Female-dark-High dose | Saline vs. PTH | 0.187078 | Unpaired *t*-test |
| Figure 4G | Male-Pretest | Saline vs. PTH | 0.951750056 | Two-way ANOVA, interaction |
|  | Male-Phase1 | Saline vs. PTH | 0.600156075 | Two-way ANOVA, interaction |
|  | Male-Phase2 | Saline vs. PTH | 1.73077E-06 | Two-way ANOVA, interaction |
|  | Female-Pretest | Saline vs. PTH | 1 | Two-way ANOVA, interaction |
|  | Female-Phase1 | Saline vs. PTH | 1 | Two-way ANOVA, interaction |
|  | Female-Phase2 | Saline vs. PTH | 0.9504759 | Two-way ANOVA, interaction |
| Figure 4F | Male-light-Pretest | Saline vs. PTH | 0.993981 | Unpaired *t*-test |
|  | Male-light-Low dose | Saline vs. PTH | 0.216833 | Unpaired *t*-test |
|  | Male-light-High dose | Saline vs. PTH | 0.483194 | Unpaired *t*-test |
|  | Female-light-Pretest | Saline vs. PTH | 0.693118 | Unpaired *t*-test |
|  | Female-light-Low dose | Saline vs. PTH | 0.827206 | Unpaired *t*-test |
|  | Female-light-High dose | Saline vs. PTH | 0.777596 | Unpaired *t*-test |
|  | Male-dark-Pretest | Saline vs. PTH | 0.252632 | Unpaired *t*-test |
|  | Male-dark-Low dose | Saline vs. PTH | 0.517977 | Unpaired *t*-test |
|  | Male-dark-High dose | Saline vs. PTH | 0.115378 | Unpaired *t*-test |
|  | Female-dark-Pretest | Saline vs. PTH | 0.829111 | Unpaired *t*-test |
|  | Female-dark-Low dose | Saline vs. PTH | 0.793299 | Unpaired *t*-test |
|  | Female-dark-High dose | Saline vs. PTH | 0.397711 | Unpaired *t*-test |
| Figure 5E | Central time | mCherry vs. hM3Dq | 0.26 | Unpaired *t*-test |
|  | Central entries | mCherry vs. hM3Dq | 0.067 | Unpaired *t*-test |
|  | Distance | mCherry vs. hM3Dq | 0.139 | Unpaired *t*-test |
| Figure 5F | Open arm time | mCherry vs. hM3Dq | 0.53 | Unpaired *t*-test |
|  | Open arm entries | mCherry vs. hM3Dq | 0.42 | Unpaired *t*-test |

## References

1. Lourida, I., et al., Parathyroid hormone, cognitive function and dementia: a systematic review. PLoS One, 2015. 10(5): p. e0127574.

2. Joborn, C., et al., Psychiatric symptomatology in patients with primary hyperparathyroidism. Ups J Med Sci, 1986. 91(1): p. 77–87.

3. Repplinger, D., et al., Neurocognitive dysfunction: a predictor of parathyroid hyperplasia. Surgery, 2009. 146(6): p. 1138–43.

4. Walker, M.D., et al., Neuropsychological features in primary hyperparathyroidism: a prospective study. J Clin Endocrinol Metab, 2009. 94(6): p. 1951–8.

5. Solomon, B.L., M. Schaaf, and R.C. Smallridge, Psychologic symptoms before and after parathyroid surgery. Am J Med, 1994. 96(2): p. 101–6.

6. Chan, A.K., et al., Clinical manifestations of primary hyperparathyroidism before and after parathyroidectomy. A case-control study. Annals of Surgery, 1995. 222(3): p. 402–414.

7. Roman, S.A., et al., Parathyroidectomy improves neurocognitive deficits in patients with primary hyperparathyroidism. Surgery, 2005. 138(6): p. 1121–8; discussion 1128-9.

8. Chiba, Y., et al., Marked improvement of psychiatric symptoms after parathyroidectomy in elderly primary hyperparathyroidism. Endocr J, 2007. 54(3): p. 379–83.

9. Espiritu, R.P., et al., Depression in primary hyperparathyroidism: prevalence and benefit of surgery. J Clin Endocrinol Metab, 2011. 96(11): p. E1737–45.

10. Roman, S.A., et al., The effects of serum calcium and parathyroid hormone changes on psychological and cognitive function in patients undergoing parathyroidectomy for primary hyperparathyroidism. Ann Surg, 2011. 253(1): p. 131–7.

11. Babinska, D., et al., Evaluation of selected cognitive functions before and after surgery for primary hyperparathyroidism. Langenbecks Arch Surg, 2012. 397(5): p. 825–31.

12. Gilli, P. and P. De Bastiani, Cognitive function and regular dialysis treatment. Clin Nephrol, 1983. 19(4): p. 188–92.

13. Driessen, M., et al., Secondary hyperparathyroidism and depression in chronic renal failure. Nephron, 1995. 70(3): p. 334–9.

14. Cengiz, K. and A. Ozkan, Depression and secondary hyperparathyroidism in chronic renal failure. Nephron, 1998. 79(4): p. 508–9.

15. Chou, F.F., et al., Cognitive changes after parathyroidectomy in patients with secondary hyperparathyroidism. Surgery, 2008. 143(4): p. 526–32.

16. Bossola, M., et al., Correlates of symptoms of depression and anxiety in chronic hemodialysis patients. Gen Hosp Psychiatry, 2010. 32(2): p. 125–31.

17. Cheng, S.-P., et al., Parathyroidectomy improves symptomatology and quality of life in patients with secondary hyperparathyroidism. 2014. 155(2): p. 320–328.

18. Harvey, S. and S. Hayer, Parathyroid hormone binding sites in the brain. Peptides, 1993. 14(6): p. 1187–91.

19. Matsui, H., et al., Central actions of parathyroid hormone on blood calcium and hypothalamic neuronal activity in the rat. American Journal of Physiology - Regulatory Integrative and Comparative Physiology, 1995. 268(1 37–1).

20. Filipovic, N., et al., Expression of PTHrP and PTH/PTHrP receptor 1 in the superior cervical ganglia of rats. Neuropeptides, 2014. 48(6): p. 353–9.

21. Usdin, T.B., C. Gruber, and T.I. Bonner, Identification and Functional Expression of a Receptor Selectively Recognizing Parathyroid Hormone, the PTH2 Receptor (&#x2217;). Journal of Biological Chemistry, 1995. 270(26): p. 15455–15458.

22. Dimitrov, E.L., et al., Neuropathic and inflammatory pain are modulated by tuberoinfundibular peptide of 39 residues. Proc Natl Acad Sci U S A, 2013. 110(32): p. 13156–61.

23. Dimitrov, E.L., E. Petrus, and T.B. Usdin, Tuberoinfundibular peptide of 39 residues (TIP39) signaling modulates acute and tonic nociception. Exp Neurol, 2010. 226(1): p. 68–83.

24. Fegley, D., et al., Increased fear- and stress-related anxiety-like behavior in mice lacking tuberoinfundibular peptide of 39 residues. Genes, Brain and Behavior, 2008. 7(8): p. 933–942.

25. Tsuda, M.C., et al., Incubation of Fear Is Regulated by TIP39 Peptide Signaling in the Medial Nucleus of the Amygdala. J Neurosci, 2015. 35(35): p. 12152–61.

26. Saggese, G., et al., Immunoreactive Parathyroid Hormone and Calcitonin in Children’s Cerebrospinal Fluid. Hormone Research in Paediatrics, 1986. 23(3): p. 177–180.

27. Johansson, P., et al., Cerebrospinal fluid (CSF) 25-hydroxyvitamin D concentration and CSF acetylcholinesterase activity are reduced in patients with Alzheimer’s disease. 2013. 8(11): p. e81989.

28. 28. Gennari, C., Parathyroid Hormone and Pain, in New Actions of Parathyroid Hormone, S.G. Massry and T. Fujita, Editors. 1989, Springer US: Boston, MA. p. 335–345.

29. Joborn, C., et al., Cerebrospinal fluid calcium, parathyroid hormone, and monoamine and purine metabolites and the blood-brain barrier function in primary hyperparathyroidism. Psychoneuroendocrinology, 1991. 16(4): p. 311–322.

30. Zhang, L., et al., Bidirectional control of parathyroid hormone and bone mass by subfornical organ. Neuron, 2023. 111(12): p. 1914–1932.e6.

31. Ni, L.-H., et al., A rat model of SHPT with bone abnormalities in CKD induced by adenine and a high phosphorus diet. 2018. 498(3): p. 654–659.

32. Chari, T., et al., The Stage of the Estrus Cycle Is Critical for Interpretation of Female Mouse Social Interaction Behavior. Frontiers in Behavioral Neuroscience, 2020. 14.

33. Tucker, L.B. and J.T. McCabe, Behavior of Male and Female C57BL/6J Mice Is More Consistent with Repeated Trials in the Elevated Zero Maze than in the Elevated Plus Maze. Front Behav Neurosci, 2017. 11: p. 13.

34. Koonce, C.J., A.A. Walf, and C.A. Frye, *Type 1 5*α*-reductase may be required for estrous cycle changes in affective behaviors of female mice*. Behavioural Brain Research, 2012. 226(2): p. 376–380.

35. Huang, K., et al., A hierarchical 3D-motion learning framework for animal spontaneous behavior mapping. 2021. 12(1): p. 2784.

36. Fang, S., et al., Sexually dimorphic control of affective state processing and empathic behaviors. 2024. 112(9): p. 1498–1517. e8.

37. Thomas, S.S. and W.E. Mitch, Parathyroid hormone stimulates adipose tissue browning: a pathway to muscle wasting. Curr Opin Clin Nutr Metab Care, 2017. 20(3): p. 153–157.

38. Hoare, S.R.J., T.I. Bonner, and T.B. Usdin, Comparison of Rat and Human Parathyroid Hormone 2 (PTH2) Receptor Activation: PTH Is a Low Potency Partial Agonist at the Rat PTH2 Receptor*. Endocrinology, 1999. 140(10): p. 4419–4425.

39. Goold, C.P., T.B. Usdin, and S.R.J. Hoare, Regions in Rat and Human Parathyroid Hormone (PTH) 2 Receptors Controlling Receptor Interaction with PTH and with Antagonist Ligands. The Journal of Pharmacology and Experimental Therapeutics, 2001. 299(2): p. 678–690.

40. Yao, Z., et al., A high-resolution transcriptomic and spatial atlas of cell types in the whole mouse brain. Nature, 2023. 624(7991): p. 317–332.

41. Nolte, E.D., K.A. Nolte, and S.S. Yan, Anxiety and task performance changes in an aging mouse model. Biochemical and Biophysical Research Communications, 2019. 514(1): p. 246–251.

42. Weber, T., et al., Parathyroidectomy, elevated depression scores, and suicidal ideation in patients with primary hyperparathyroidism: results of a prospective multicenter study. 2013. 148(2): p. 109–115.

43. Weber, T., et al., Symptoms of primary hyperparathyroidism in men and women: the same but different*?* 2020. 36(1): p. 41–47.

44. Natsui, K., et al., A high abundance of PTH-related protein in human cerebrospinal fluid. Journal of Bone and Mineral Metabolism, 1994. 12(1): p. S207–S209.

45. Liu, Y., et al., An optogenetic approach for regulating human parathyroid hormone secretion. 2022. 13(1): p. 771.

46. Ni, L.-H., et al., A rat model of SHPT with bone abnormalities in CKD induced by adenine and a high phosphorus diet. Biochemical and Biophysical Research Communications, 2018. 498(3): p. 654–659.

